# Deconer: A comprehensive and systematic evaluation toolkit for reference-based cell type deconvolution algorithms using gene expression data

**DOI:** 10.1101/2023.12.24.573278

**Authors:** Wei Zhang, Xianglin Zhang, Qiao Liu, Lei Wei, Xu Qiao, Rui Gao, Zhiping Liu, Xiaowo Wang

## Abstract

In recent years, computational methods for quantifying cell type proportions from transcription data have gained significant attention, particularly those reference-based methods which have demonstrated high accuracy. However, there is currently a lack of comprehensive evaluation and guidance for available reference-based deconvolution methods in cell proportion deconvolution analysis. In this study, we propose a comprehensive evaluation toolkit, called Deconer, specifically designed for reference-based deconvolution methods. Deconer provides various simulated and real gene expression datasets, including both bulk and single-cell sequencing data, and offers multiple visualization interfaces. By utilizing Deconer, we conducted systematic comparisons of 14 reference-based deconvolution methods from different perspectives, including method robustness, accuracy in deconvolving rare components, signature gene selection, and building external reference. We also performed an in-depth analysis of the application scenarios and challenges in cell proportion deconvolution methods. Finally, we provided constructive suggestions for users in selecting and developing cell proportion deconvolution algorithms. This work presents novel insights to researchers, assisting them in choosing appropriate toolkits, applying solutions in clinical contexts, and advancing the development of deconvolution tools tailored to gene expression data.

## Introduction

The rapid development of high-throughput sequencing technologies, such as RNA-seq and scRNA-seq, provide us unprecedented opportunities to unravel the global gene expression in cells [1,2]. In multicellular organisms, nearly all physiological and pathological processes involve multiple types of cells, each playing a unique role in specific mechanisms [3,4]. Quantifying cell proportions is essential for tracking signals associated with certain phenotypes or diseases. For example, predicting cell proportion aids in uncovering the cellular composition and heterogeneity of tissues and organs, providing vital information for studying cell systems [5]. Estimation of cell type proportions helps researchers understand the infiltration of immune cells in the tumor microenvironment, which is crucial for selecting appropriate immunotherapy strategies [6]. Additionally, by analyzing the proportion of cell types in tissue samples, we can enhance the accuracy of disease diagnosis and determine the stage of disease progression [7,8].

So far, the proportions of different cell types can be determined through various experimental methods based on physical properties, such as flow cytometry, immunohistochemistry and fluorescence-activated cell sorting (FACS) [9,10]. However, these techniques have limitations in certain aspects. For instance, flow cytometry is the gold-standard for identification of cell subsets, but its throughput might be relatively low [11–13].

Furthermore, the application of FACS relies on the availability of suitable cell surface markers. In some cases, the lack of specific markers may result in unfeasibility to effectively capture and identify particular cell types [12,14]. The advance of single-cell sequencing technologies has revolutionized our understanding of cellular heterogeneity and allowed us to probe the functional role of each cell type within complex multicellular systems. However, a considerable number of bulk samples are still generated in clinical and experimental researches and numerous bulk gene expression data in existing databases remain underutilized and untapped. Therefore, computational models for quantifying cell type proportion remain highly relevant and essential.

The proportions of different cell types can be inferred from various types of omics data, including gene expression data (microarray, RNA-seq), DNA methylation data (WGBS, RRBS), as well as DNA accessibility data (ATAC-seq) [15–18]. Currently, DNA accessibility data was utilized to infer cell proportions in some studies, but there is limited research in this regard. DNA methylation data is one of the most commonly used data types for this task. While providing valuable insights into gene regulation, this type of data involves a more complex experimental and data processing procedures, which indicates it is more suitable for assisting in the prediction of cell proportion when analyzing transcriptional regulation. Of course, in certain situations, DNA methylation data may be more appropriate for deciphering cell proportions, such as predicting circulating tumor DNA burden in cell-free DNA samples [7,19,20]. Gene expression data, on the other hand, directly reflects transcriptional differences among various cell types. These differences can serve as features for identifying different cell type proportions. Furthermore, gene expression data has a direct connection to cellular functions and biological processes, making cell proportion deconvolution results more biologically meaningful. Compared to the previously mentioned data types, gene expression data can be more easily obtained experimentally and can be processed using standard normalization methods, such as FPKM (Fragments Per Kilobase of transcript per Million mapped reads) and TPM (Transcripts Per Million).

Cell type deconvolution algorithms for gene expression data can be primarily divided into two categories: reference-based methods and reference-free methods. Reference-based methods utilize pre-defined cell type-specific gene expression profiles as prior information and apply techniques such as constrained linear regression to estimate cell proportions. In contrast, reference-free methods do not rely on external expression references and directly obtain cell proportions and expression profiles using techniques such as non-negative matrix factorization (NMF). Reference-free methods need to simultaneously estimate expression profiles and cell proportions, which might attenuate the deconvolution performance when compared with reference-based methods [18,21–24]. Hence, it is crucial to evaluate and compare these reference-based deconvolution methods for swiftly and accurately deciphering the proportions of distinct cell types.

In this study, we developed a dedicated tool called Deconer (<u>Decon</u>volution <u>E</u>valuator, Figure 1) to facilitate the systematic comparisons. Deconer incorporates numerous simulation data generation methods based on both bulk and single-cell gene expression data, as well as a wide range of evaluation metrics and visualization tools to aid researchers in their analysis. In addition, we collected a large set of high-quality RNA-seq and scRNA-seq data that can be used for evaluating the available deconvolution methods. Some of these datasets are experimentally obtained with known cell proportions, providing valuable resources for method comparison. We also conducted a systematic evaluation analysis of 14 reference-based deconvolution algorithms for estimating cell type proportions, including 12 conventional deconvolution methods based on either probabilistic models or machine learning models, as well as two recently proposed deep learning-based methods (Table 1). Factors that influence deconvolution procedures like data noise, the number of cellular components as well as the presence of rare components were also taken into consideration. Multiple evaluation metrics were used for analyzing and interpreting the results, providing guidance on selecting suitable deconvolution methods for gene expression data in different research scenarios. All analyses in this assessment is performed using Deconer, facilitating readers to reproduce the results and enabling further comparison and evaluation of related deconvolution works in the future. We have made Deconer and the processed datasets publicly available, allowing for easier result reproduction and comparison of new deconvolution algorithms in future.

**Figure 1.**
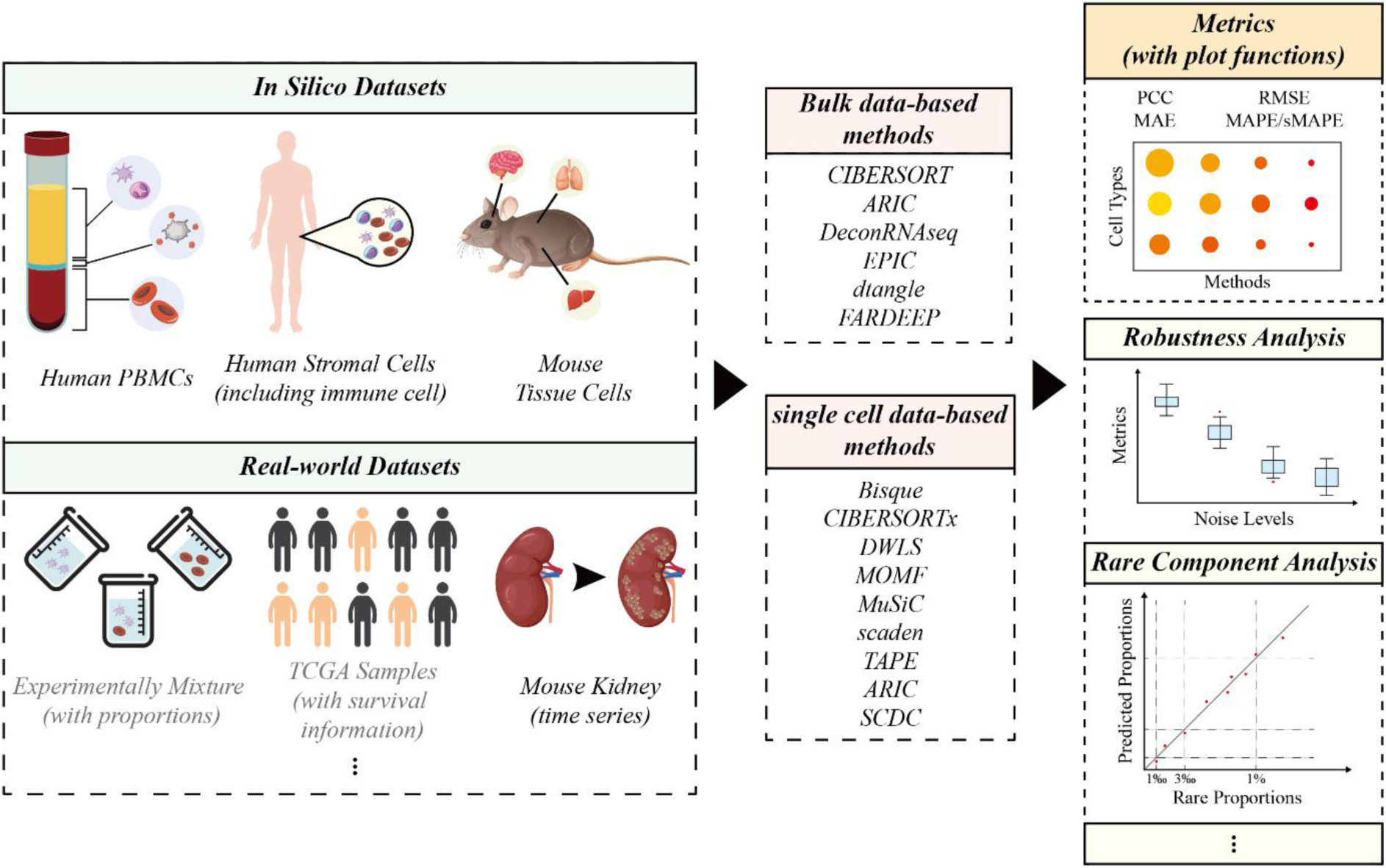
The overview workflow of this study and the main functions in Deconer. The left panel shows the datasets provided by Deconer. The data marked in black font were utilized in this study, while the data provided by Deconer in gray font were pre-processed but not used in the comparison study conducted in this study. The middle panel lists all the deconvolution methods. Some main functions provided by Deconer is shown in the right panel.

**Table 1.** The deconvolution algorithms for gene expression data involved in this study.

| Deconvolution methods | Reference data type | Description and remark |
| --- | --- | --- |
| ARIC[4] | Bulk & single cell | The method adopts component-wise condition number and can be used for multiple data types. Implemented as a python package (version 0.1.0). |
| Bisque[40] | single cell | Non-negative least-squares (NNLS) regression is used for estimating cell proportions. Implemented as an R package (version 1.0.5). |
| CIBERSORT[39] * | Bulk | A well-known deconvolution method adopts linear v-SVR. |
| CIBERSORTx[10] | single cell | The update version of CIBERSORT with user-friendly web interface. |
| DeconRNAseq[36] | Bulk | One of the earliest deconvolution methods using constrained linear regression model. Implemented as an R package (version 1.40.0). |
| dtangle[3] | Bulk | A newly proposed method which is robust to outliers. Implemented as an R package (version 2.0.9). |
| DWLS[41] | single cell | This method adopts dampened weighted least squares is used and can correct common biases towards cell types. Implemented as an R package (version 0.1.0). |
| EPIC[37] | Bulk | This method is robust to unknown mixture content. Implemented as an R package (version 1.1.6). |
| FARDEEP[38] | Bulk | FARDEEP adopts least trimmed squares and is robust to outliers. Implemented as an R package (version 1.0.1). |
| MOMF[42] | single cell | MOMF is based on poisson distribution and alternating direction method of multipliers. Implemented as an R package (version 0.2.0). |
| MuSiC[43] | single cell | MuSiC assigns weights to genes to maintain cross-subject and cross-cell consistency. Implemented as an R package (version 1.0.0). |
| scaden[44] | single cell | The first proposed deep learning-based deconvolution algorithm. Implemented as a python package (version 1.1.2). |
| SCDC[46] | single cell | A method adopts an ENSEMBLE strategy for addressing the problem of batch-effect confounding. Implemented as an R package (version 0.0.0.9000). |
| TAPE[45] | single cell | A method based on auto-encoder architecture. Implemented as a python package (version 1.1.2). |

\* We used the version implemented in EpiDISH package (version 2.14.1) since the website of CIBERSORT is archived.

## Methods

### Toolkits

We developed Deconer, a comprehensive evaluation toolkit for cell type deconvolution algorithms, as a free R package. The main functions of Deconer are displayed in Figure 1. Deconer includes a variety of functions for performance assessment. The aforementioned pseudo bulk data can be directly generated through Deconer. Deconer also offers a diverse range of preprocessed real experimental datasets with known cell proportions [25–32]. A gene expression dataset of experimental simulation of chronic kidney disease in mouse kidney is also provided. In addition, the gene expression data from 32 TCGA cancer types along with patients’ survival information are also included [33–35]. Deconer integrates a variety of evaluation metrics and plotting programs. Furthermore, it offers several evaluation functions, such as stability testing of the model under simulated noise conditions, and accuracy analysis of rare component deconvolution.

### Overview of reference-based cell type deconvolution algorithms for gene expression data

Reference-based deconvolution methods can be mainly classified into two subcategories: those using bulk data as a reference and those using single-cell data as a reference. Typical methods employing bulk data as a reference include DeconRNAseq [36], EPIC [37], dtangle [3], FARDEEP [38] and CIBERSORT [39], while those utilizing single-cell data as a reference encompass Bisque [40], CIBERSORTx [10], DWLS [41], MOMF [42], MuSiC [43], scaden [44], TAPE [45] and SCDC [46]. Notably, both bulk and single-cell data can serve as external reference for the ARIC [4]. Other methods that have already been compared and analyzed in previous studies are not included in our current analysis [18,23,37]. Among these methods, the majority are based on the linear regression model expect for scaden and TAPE, where the measured sample gene expression values are a summation of gene expression values from different types of cells. Although these linear regression-based methods share a common conceptual foundation, they differ in focal points and approaches to proportional estimation. For example, DeconRNAseq adopts quadratic programming for estimating the mixing proportions of distinctive cell types while Bisque utilizes non-negative least-squares [36]. EPIC aims to achieve precise differentiation and proportion estimation of distinct cell types by seeking out genes with highly specific expression [37]. dtangle uses a biologically appropriate linear mixing model and using log-transformed data to fit the model [3]. CIBERSORT estimates relative subsets of RNA transcripts using SVR and CIBERSORTx is an update version for single cell data [10,39]. MOMF directly models gene count data using a Poisson distribution and employs the Alternating Direction Method of Multipliers (ADMM) approach to reduce computational load [42]. FARDEEP, ARIC, DWLS as well as MuSiC achieve the auto-selection or weighting of different signature genes by analyzing the expression profiles of various cell types and employ approaches such as adaptive deconvolution to estimate proportions accurately [4,38,41,43]. SCDC, on the other hand, employs an ensemble learning approach, integrating multiple datasets for deconvolution to enhance model stability [46]. It is worth mentioning that this study incorporates two recently proposed deconvolution methods based on deep learning techniques, namely scaden and TAPE. scaden is the first deconvolution method to employ deep learning techniques [44]. It employs a multi-layer fully connected neural network that directly takes high variance genes as input to estimate proportions of different cell types. On the other hand, TAPE utilizes an auto-encoder architecture, simultaneously estimating both cell type proportions and gene expression profiles [45]. All the above-mentioned methods and the corresponding descriptions are summarized in Table 1.

### Evaluation datasets

In this paper, we extended the former collected data gathered in our previous work with additional bulk and single-cell RNA-seq datasets [4]. In total, we provided four types of pseudo-bulk data; two were generated through simulation of bulk RNA-seq data, while the other two were generated through simulation of scRNA-seq data. Inspired by the Tumor Deconvolution DREAM Challenge, we collected 233 high-quality bulk RNA-seq samples to simulate cell populations, including stromal cells and immune cells. We constructed two distinct scenarios, corresponding to “coarse” and “fine” conditions respectively. Under the coarse condition, mixed samples will contain 8 different cell types, whereas under the fine condition, mixed samples will contain 14 different cell types. Additionally, we collected a unique single-cell dataset consisting of seven different tissues from fetal mouse [47]. In this dataset, each cell type has 1,500 single-cell data entries, which ensures adequate data for the construction of simulation sample and external references. Furthermore, given the relatively large expression profile differences among these seven tissues, samples generated using this dataset can provide a fair performance evaluation for various algorithms [47]. We also collected 10K human PBMC scRNA-seq data. For this dataset, we employed MUON to annotate 13 reliable cell types of 11094 cells (see Supplementary Figure S1) [48].

In addition to the simulation data, we also employed a well-characterized mouse kidney dataset to test the comparing methods [33,34]. This dataset represents a real-world case with unknown proportions, and we assessed the accuracy of the derived cell proportions by associating them with known biological phenotypes or processes.

We adopted different approaches for generating evaluation data from bulk or single cell gene expression data. For bulk data, we first divided all samples into training (70%) and test sets (30%). The training data was solely used to generate the external reference, while the test data was utilized to produce samples for assessment. Evaluation samples were directly generated using the same linear mixing approach as ARIC [4]. In brief, numerical linear combinations were performed after generating cell proportions according to the requirements of different evaluation criteria. As for scRNA-seq data, we initially divided the cells for each cell type into a training set (70%) and a test set (30%) based on the number of cells. Training set was also only used for generating external reference. To generate pseudo bulk data, we first created random cell proportions and then sampled cells from the test set based on the required cell number. Then, the gene expression values of cells were added together to simulate the bulk sample. More detailed descriptions of the data generation methods under different scenarios are provided in each section.

The preprocessing workflow for all datasets is provided in the Supplementary Materials Section 2, and all the collected data are summarized in Table 2, Supplementary Table S1 and S2. All datasets are provided in Deconer website (https://honchkrow.github.io/Deconer_dataset/).

**Table 2.** Summary of the test datasets used in this study.

| Dataset name | Number of<br>cell types | Data type | Description |
| --- | --- | --- | --- |
| coarse_bulk | 8 | Bulk | This dataset is from 233 high-quality bulk RNA-seq samples to simulate stromal cell populations, including immune cell populations. For each cell type, RNA-seq samples are collected from at least two distinct studies. |
| fine_bulk | 14 | Bulk | This dataset, like the coarse dataset, has data for all cell types derived from at least two different datasets, except for regulatory T cells, which are only sourced from a single dataset. |
| mouse_tissue | 7 | Single cell | This dataset contains 7 well-characterized mouse tissues. |
| human_PBMC | 13 | Single cell | 10k human PBMC from 10X. |
| mouse_kidney | 13 | Single cell | This data is from unilateral ureteric obstruction mouse model of renal fibrosis in kidney diseases. |

Deconer provides all of the datasets in this table, and it also offers additional datasets with known cell proportions. For detailed summary table, please see https://honchkrow.github.io/Deconer_dataset/.

### Simulation of data noise for gene expression data

According to the study by Jin et al., the negative binomial model was found to be highly effective in capturing salient features in real data, such as mean-variance trends and sample-sample concordance [18]. The model offered a depiction of the noise structure intrinsic to real data, thereby enabling a more accurate interpretation and analysis. Inspired by these findings, we adopted the negative binomial model to simulate various levels of noise as the previous work [18,49]. In brief, the model can be described as follows.

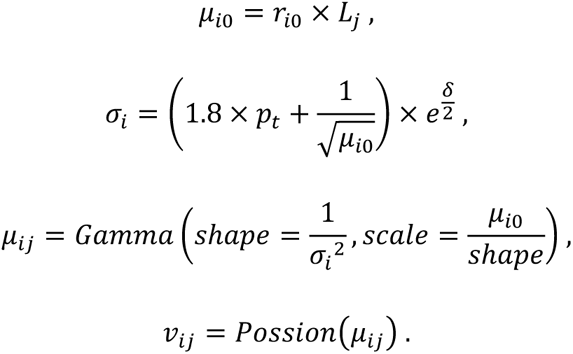

In the above equations, *δ*∼*N*(0,0.25). *r*_*i*0_ is the expected count for gene *i* in a cell type and *L*_*j*_ is the library size of sample *j*. Therefore, *μ*_*i*0_ is the expected count in the sample to be deconvoluted. We set *p*_*t*_ in a range from 0.1 to 1 to control noise level. Noise is added using *gamma* and *possion* distribution [18].

### Evaluation metrics

In this study, we utilized multiple assessment metrics in a comprehensive manner to enable an objective evaluation of various deconvolution algorithms from different perspectives. The Pearson Correlation Coefficient (PCC) was used as a fundamental measure to assess the linear relationship between the actual cell proportions and their respective predictions with different deconvolution methods. We also utilized the Mean Absolute Percentage Error (MAPE) as well as Symmetric Mean Absolute Percentage Error (sMAPE) to evaluate the degree of deviation in the prediction of each cell type. The definition of PCC, RMSE, MAPE as well as sMAPE are as follows

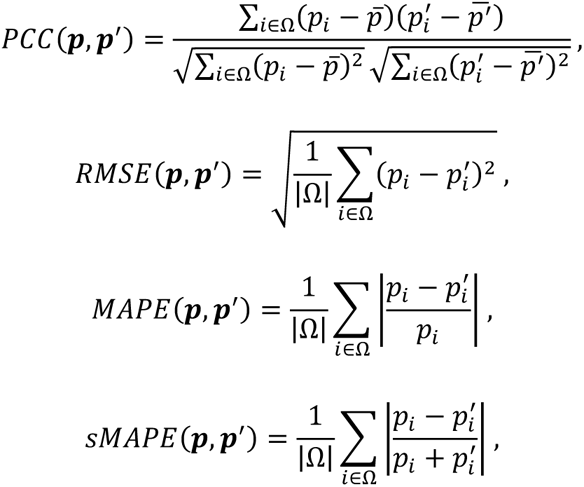

where *p*_*i*_ and 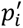 denotes the ground truth and the predicted proportion value respectively. It is noteworthy that the aforementioned metrics in Deconer can be applied to various evaluation scenarios, including the assessment of all cell types in individual sample or across multiple samples, as well as individual evaluations of different cell types across multiple samples. Accordingly, the term Ω is used to denote the cell type set in each condition and |Ω| is the number of cell types.

## Results

### Comparison of deconvolution algorithms for gene expression data under diverse levels of noise

When processing gene expression data, it is imperative to address various forms of data noise. For instance, the same cell type under different physiological states may exhibit varied gene expression patterns, thus introducing a biological noise that is difficult to be avoided. In addition, technical noise also poses a substantial challenge. This kind of noise originates from discrepancies and biases in experimental operations and data pre-processing, such as library construction and sequence alignment [50]. Both biological and technical noise can profoundly impact the cell type deconvolution from gene expression data, potentially leading to skewed or inaccurate results.

We conducted an extensive evaluation of deconvolution algorithms for bulk and single-cell gene expression data. This assessment focused on the stability of deconvolution under varying degrees of noise. For the experiments, we generated 50 samples under each noise condition and computed different metrics for each cell type. This comprehensive evaluation approach allows us to understand how these deconvolution methods perform under different levels of noise, which is a critical aspect considering the inherent noise found in real data. By generating multiple samples under various noise conditions, we can gain a holistic understanding of the robustness and reliability of these deconvolution algorithms for detecting cell types from gene expression profiling data.

As shown in Figure 2A, 2B as well as Supplementary Figure S2 and S3, it is evident that as data noise increase, there is a corresponding decline in the performance across various deconvolution algorithms. Algorithms such as ARIC, FARDEEP, DWLS, and MuSiC demonstrate superior performances under conditions without noise. However, as noise levels intensify, these algorithms exhibit a significant increase in RMSE and a drop in PCC, consistent with our expectations. The stability of DeconRNAseq, EPIC, Bisque, and MOMF is relatively lower in comparisons. Among the selected algorithms, DWLS and scaden exhibit remarkable robustness in scenarios with different levels of noise, maintaining consistently low levels of RMSE as shown in Figure 2B. This indicates that DWLS and scaden comprehensively address and discriminate all cellular components. In addition, the MuSiC and SCDC algorithms also showcase impressive deconvolution performance, standing out among the evaluated methods.

**Figure 2.**
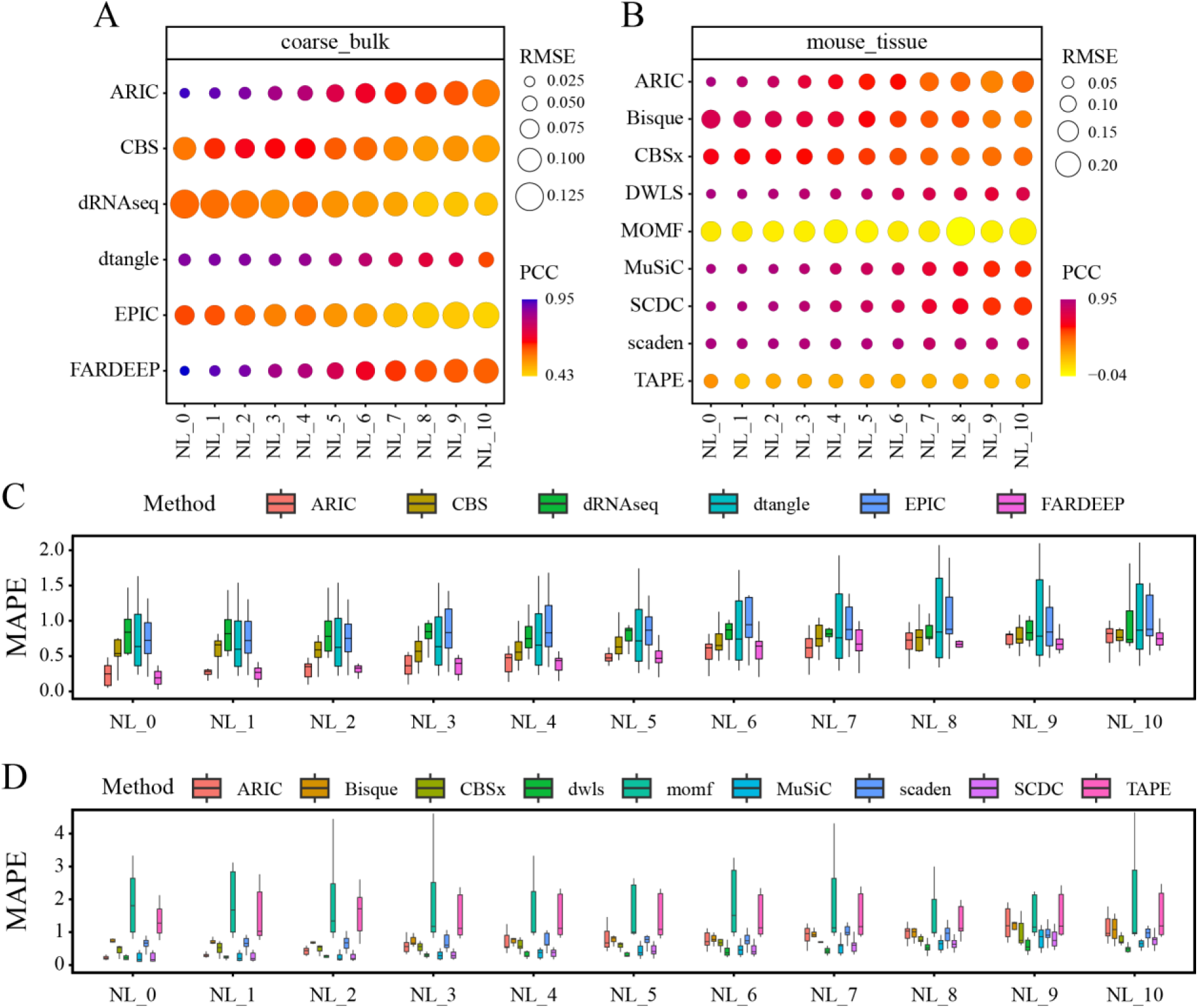
Stability testing of various deconvolution methods in scenarios with few cell types. (A) and (B) are the RMSE and PCC for methods employing bulk data and single cell data as a reference respectively. (C) and (D) are the MAPE for methods employing bulk data and single cell data as a reference respectively. We set *p*_*t*_ in a range from 0.1 to 1 to control noise level (NL_1 to NL_10). NL_0 denotes the absence of noise incorporation. CBS: CIBERSORT, dRNAseq: DeconRNAseq. CBSx: CIBERSORTx.

Our analysis further revealed that despite similar performances in terms of PCC and RMSE, there was a substantial discrepancy in the MAPE among certain methods, as demonstrated in Figures 2C and 2D. This phenomenon implies the presence of cell-type bias in many methods. For instance, we generated the scatter plots of dtangle and FARDEEP at three different noise levels in Figure 3. From Figure 2, the deconvolution results of dtangle demonstrated exceptional stability. However, its predictions exhibit a pronounced cell-type bias in Figure 3. Specifically, its predictions for endothelial cells and neutrophils were consistently under-estimated, while CD4 T cells and CD8 T cells were over-estimated. Similarly, the results of FARDEEP also manifested such bias, with B cells and monocyte cells showing noticeable deviations. Due to the unique gene expression patterns of each cell type, the limitations of deconvolution methods in capturing these intricate differences may lead to deviations between the identified cell proportions and their actual values. At present, the majority of deconvolution methods resolve this issue by identifying differentially expressed genes. For instance, EPIC proposes the identification of genes specifically expressed in a single cell type [37]. However, the search for such genes is often challenging, and it may result in an insufficient number of markers for deconvolution. On the other hand, methods like scaden and TAPE directly identify high variance genes by setting a threshold, which could potentially cause an imbalance in the quantity of markers across different cell types [44,45]. Hence, the question of how to select markers to balance the accuracy of deconvolution results and the model’s attention to different cell types warrants further exploration.

**Figure 3.**
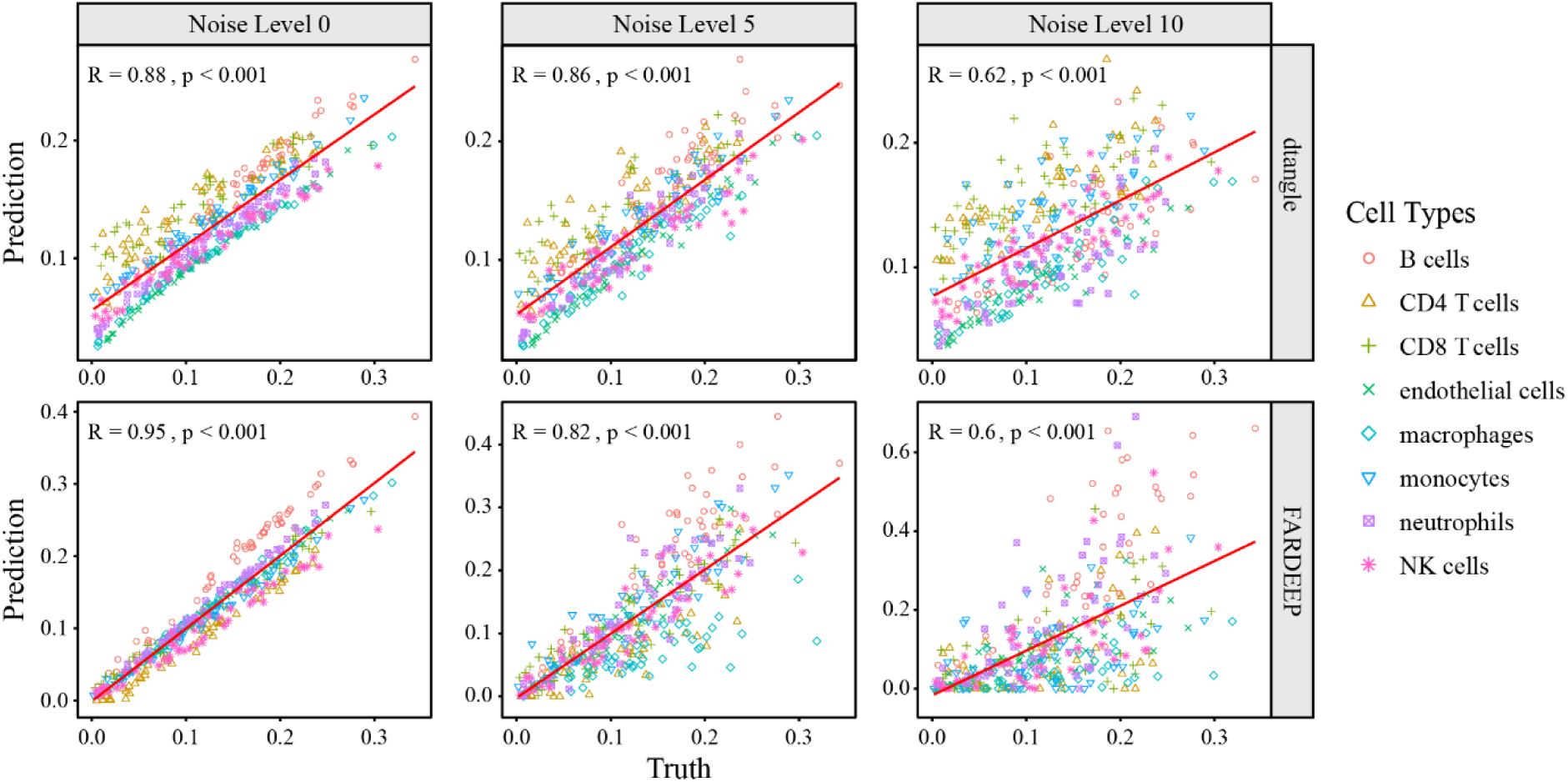
Scatter plot for dtangle and FARDEEP. PCCs and p-value are shown in top-left corner of each panel.

### Unraveling the challenge of characterizing additional cell subtypes and building external references

In the real deconvolution scenarios, the issues we encountered are substantially more complex. For example, in many research scenarios, the investigation of the proportions of subtype cells within a certain cell type is particularly significant [51–53]. Furthermore, in the absence of a high-quality external references during the deconvolution process, we can only rely on a limited number of cell expression data. These data often grapple with issues such as batch effects and different data origin, posing significant challenges to the deconvolution process [12]. These situations are prevalent in almost all deconvolution problems, and the issues of batch effects and differences in experimental origin have been extensively discussed in previous studies [12,44,53,54]. Thus, in this context, we primarily delve into a detailed discussion concerning about more cell subtypes and the difficulties encountered in constructing external reference.

This study utilizes four datasets (coarse_bulk, fine_bulk, mouse_tissue and human_PBMC) to carry out the simulation and deconvolution. The coarse_bulk and mouse_tissue datasets comprise fewer cell types, and the similarity among these cell types is relatively weak. In the contrast, fine_bulk and human_PBMC encompass a broader array of cell types, 14 and 13 respectively, with several cells being further divided into more subtypes. For instance, in the coarse_bulk dataset, we have B cells, CD4 T cells, and CD8 T cells. In the fine_bulk dataset, these three cell types are further subdivided into memory B cells, naïve B cells, memory CD4 T cells, naïve CD4 T cells, memory CD8 T cells, naïve CD8 T cells, and regulatory T cells. The functionally similar cell types share similar gene expressions, which can lead to increased cell-to-cell confusion in the deconvolution process [53,55,56]. Simultaneously, unlike the mouse_tissue dataset where each cell type has 1500 high-quality sequenced cells, the cell number of each type in the human_PBMC dataset varies significantly. This disparity renders the construction of an external reference less stable, thereby posing challenges to the deconvolution process. We deliberately escalated the difficulty of our comparison, utilizing the fine_bulk and human_PBMC datasets to assess the efficacy of the deconvolution methods.

In our investigation with the aforementioned datasets, we observed notable disparities in the performances of various deconvolution methods, as depicted in Supplementary Figure S4-S8. The methods such as CIBERSORT, CIBERSORTx, and SCDC maintained consistent performance trends as the former, demonstrating a clear decrease in deconvolution efficiency as the level of noise increased. ARIC, FARDEEP, DWLS, and MuSiC methods continued to demonstrate exceptional performance even when confronted with an increased number of cell subtypes. The stable performance might be attributed to the specific designs of these deconvolution methods. ARIC and DWLS emphasize the consideration of rare proportion components, while FARDEEP and MuSiC opt for the selection of specific features through outlier filtering or tree-based methods. These two aspects are particularly crucial for the accurate deconvolution of cell subtypes. The results of dtangle remained consistent and stable, corroborating the findings presented in Figure 2. It is noteworthy that two deep learning-based methods, scaden and TAPE, exhibited relatively stable performances. Both scaden and TAPE are built upon deep neural networks, which are exemplary model-free machine learning techniques with strong adaptability to data [44,45,57]. These methodologies leverage single-cell data to generate numerous mixed samples for model training, which guarantees that they are capable of maintaining their performance, even when faced with more complex data and special conditions, highlighting their potential utility in the context of cell deconvolution from gene expression data.

However, in the human_PBMC dataset, the predictive accuracy for the proportion of certain individual cell types, such as pDC and mDC, is substantially compromised (see Supplementary Figure S8). These cell types are represented in very low quantities, presenting considerable challenges in effectively simulating their proportional presence under complex conditions. This scarcity of specific cell types in the dataset contributes to a decreased prediction accuracy. Therefore, it is crucial for a deep learning-based methods to ensure the availability of a sufficient number of high-quality single cells for the generation of simulated samples in training. The quality of the single cell data directly influences the learning process, where the representativeness and diversity of the data are paramount for the model to understand complex patterns and make accurate predictions. The utilization of a larger set of high-quality single cells increases the robustness of the training procedure, thereby enhancing the overall reliability and predictive power of the model. This data enrichment allows for better generalization when the model encounters complex situations, such as predicting the proportion of specific cell types in a given sample. Therefore, the usage of a sufficient number of high-quality single cells is not just beneficial but essential for training a deep learning model to ensure its reliability and validity in diverse and complex scenarios.

Besides, we found that the issue of cell type proportion prediction bias almost invariably observed in all methods (see Supplementary Figure S7 and S8). Among these methods, ARIC, FARDEEP, DWLS and MuSiC, which inherently incorporate gene feature selection mechanisms, demonstrated a superior deconvolution performance. On the contrary, other methods that do not natively incorporate feature selection exhibited the bias more conspicuously (see Supplementary Figure S7 and S8). scaden and TAPE, which directly use highly variable genes, had even more pronounced cell type biases. Hence, when estimating proportions for finer cell subtype, the selective pick of marker genes is imperative, otherwise, biases stemming from cell type similarities are extremely likely to occur.

### Deconvolution of rare components

Accurately estimating the proportions of specific cell types, especially rare cell types in certain circumstances, is of crucial importance in uncovering their significance in specific phenotypes or diseases [4,20]. For instance, precise estimation of the proportion for tumor-infiltrating lymphocytes (TILs) plays a vital role in predicting clinical prognosis and developing personalized treatment strategies [58–60]. TILs are lymphocytes present in tumor tissues and are critical components of the immune system. They interact with tumor cells, regulating and influencing tumor growth, progression, and treatment response. TILs encompass various immune cell subtypes, including CD8+ T cells, CD4+ T cells, and natural killer (NK) cells [61,62]. However, studies have revealed variations in the proportions of TILs among patients with the same cancer type. The presence of TILs and other immune cells in tumor tissue is referred to as a hot tumor. Hot tumors with a high proportion of TILs are known to exhibit strong immune responses, which are often associated with favorable prognosis and increased sensitivity to immunotherapy. In contrast, cold tumors lack significant TILs and other immune cells in the tumor tissue, resulting in a relatively weak immune response. Cold tumors are typically associated with poor prognosis and low sensitivity to immunotherapy. Therefore, accurate analysis of the proportions of TILs in the tumor microenvironment of different patients is crucial for understanding the immune characteristics of tumors and providing important guidance for clinical classification and treatment strategies. In addition, research on circulating free RNA (cfRNA) also requires the accurate estimation of the proportion of rare cell types [63–66]. cfRNA originates from processes such as cell apoptosis and secretion [64,66]. Due to their active division and metabolism, cancer cells release cfRNA into the blood. Certain genes known as “dark channel biomarkers” have been found to retain tissue and cancer-specific characteristics in cfRNA [67]. This discovery provides new avenues for future cancer detection, prediction of tumor tissue origin, and determination of cancer subtypes. However, in early-stage cancer patients, the size of the lesion tissue is small, resulting in an extremely low proportion of cfRNA released by cancer cells into the blood, making detection extremely challenging.

For this problem, we conducted tests on the presence of rare components using the aforementioned deconvolution methods and evaluated the sensitivity of different algorithms to rare components. Following the approach used by Song et al., we set up a gradient of proportions to simulate different scenarios of rare components [12]. The gradient proportions were set as 0.001, 0.003, 0.005, 0.008, 0.01, 0.03, and 0.05 respectively. Due to the small scale of the simulation proportions, the MAPE can yield large percentage errors for predicted values that are zero or close to zero, potentially causing biased evaluation results. Therefore, we chose to use the sMAPE to measure the relative prediction error.

In our evaluation experiments, we tested the above-mentioned methods on the four datasets provided by Deconer and compared their performance in terms of RMSE and sMAPE metrics. Overall, ARIC, FARDEEP, DWLS, and MuSiC demonstrate remarkable performance in predicting minor components (Figures 4 and Supplementary Figure S9). However, the results revealed that many methods tend to predict rare components as 0, like Bisque, EPIC, scaden and TAPE. For example, in the mouse_tissue dataset, Bisque predicted all rare components as zero (see Supplementary Figure S9). We believe that this is because many methods focus on measuring errors such as MAE, RMSE or weighted RMSE, which tend to emphasize errors in larger proportion components while neglecting predictions for rare components. Although these methods often achieve good RMSE scores, their performance is inadequate in handling rare proportion components as indicated by the sMAPE metric.

**Figure 4.**
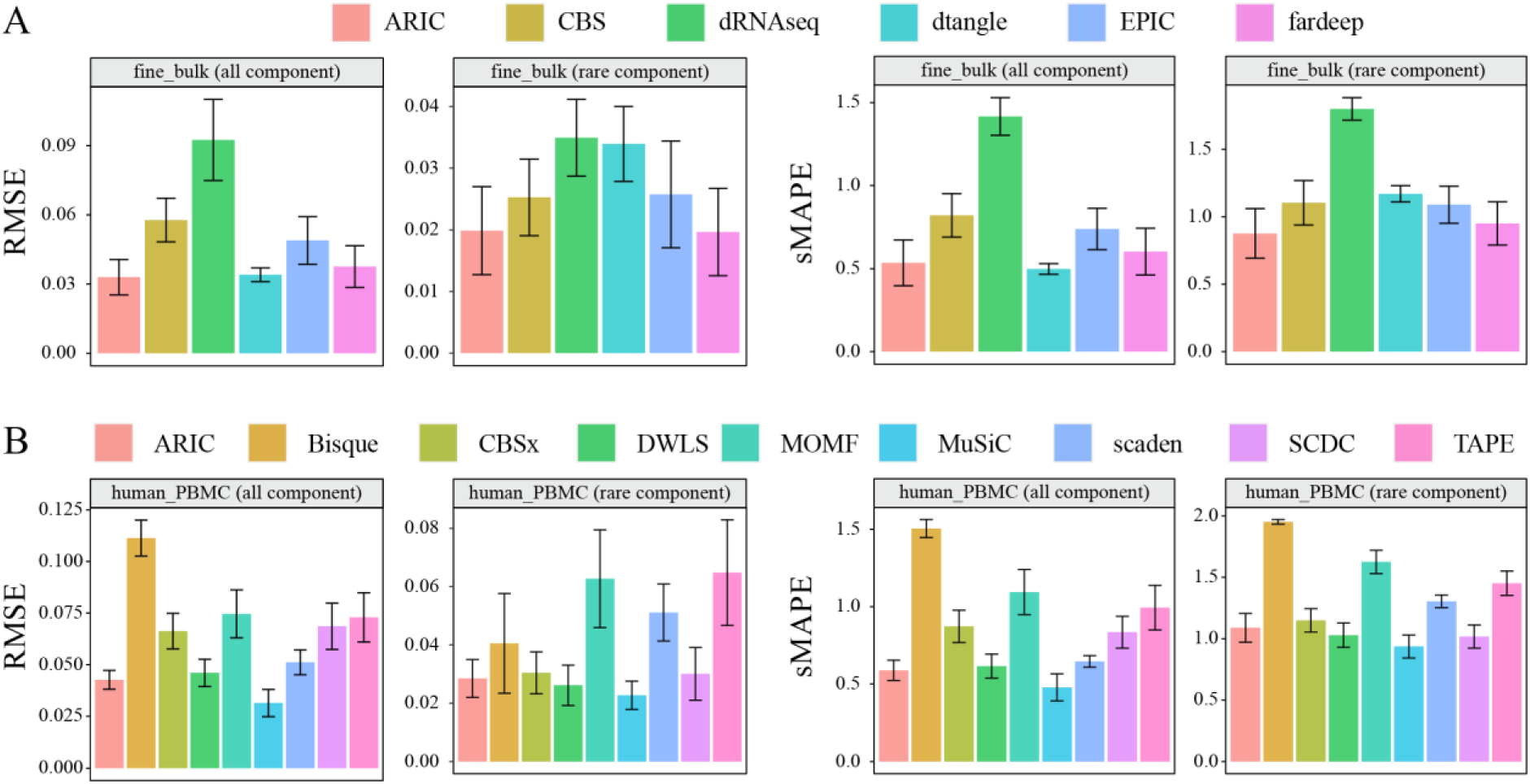
Impact of rare components on the deconvolution results using fine_bulk and human_PBMC datasets. All components as well as rare components are illustrated respectively. CBS: CIBERSORT, dRNAseq: DeconRNAseq. CBSx: CIBERSORTx.

Scaden and TAPE are two recently proposed deep learning methods that utilize random proportions for training data generation in the absence of prior knowledge [44,45]. However, these approaches face difficulties in comprehensively simulating crucial rare components. Therefore, it is preferable to perform simulation data generation based on prior information. Notably, TAPE employs the Dirichlet distribution as a tool for generating the mixture proportions in simulated data [45]. This distribution allows for flexible control of component ratios by adjusting the parameters set for the Dirichlet distribution. Users can simulate components by fine-tuning these parameters, resulting in generated data that closely aligns with real-world scenarios.

### Impact of training data generation on deep learning-based deconvolution methods

In recent years, deep learning-based deconvolution methods, especially the scaden, have received significant attention in numerous cutting-edge studies. Multiple works have demonstrated their excellent performances in cell type deconvolution across various conditions [68–70]. By harnessing the powerful feature selection capability underlying deep learning, scaden enables automatic selection of signature genes and deconvolute of expression values by simply inputting highly variable genes. Furthermore, deep learning-based methods exhibit strong robustness to input data noise and bias, surpassing the capabilities of conventional regression-based deconvolution methods [44]. Building upon scaden, TAPE has made further advancements by employing an autoencoder to reconstruct gene expression profiles, thereby enhancing the robustness [45]. Although these deep learning-based methods have achieved remarkable success, our analysis reveals the significant impacts of data generation approaches on the effectiveness of these methods.

For scaden and TAPE, there are certain differences in the way they generate training data. Specifically, scaden directly samples and normalizes from a uniform distribution, which is similar to the approach used in some other studies [4,15]. On the other hand, the TAPE method generates random proportions using the Dirichlet distribution, which can better simulate real-world scenarios when there are certain prior proportions [45]. However, in most cases, obtaining accurate prior proportions is challenging. Therefore, in general, these two data generation approaches struggle to achieve good coverage in the proportion space, especially when there are a large number of cell types, making it difficult to accurately detect cell types under extreme proportion conditions. Additionally, since the generation of training data relies on external single-cell sequencing datasets, it is important to have an adequate number of cells for each cell type and preferably maintain a certain level of consistency. However, this requirement contradicts the issue of cellular heterogeneity in single-cell data [71]. Thus, the careful selection of single-cell data for training becomes crucial.

Additionally, we found that the number of cells in mixed samples during training also has an impact on the deconvolution performance. In the scaden method, this value is set to 100, and users can adjust it according to their needs. In contrast, the TAPE method sets this value to 500 and does not support adjustment (in the latest version 1.1.2). To verify this phenomenon, we conducted tests using the mouse_tissue dataset by using the original implementation of the two methods. For scaden, we generated 50 pseudo-bulk samples, each consisting of 3000 single cells. We then adjusted the number of cells in the training samples generated by scaden, setting them to 100, 500, 1000, 2000, 3000, 4000, and 5000, respectively. As for TAPE, we generated pseudo-bulk samples with cell numbers of 100, 300, 500, 700, 1000, and 3000, and kept the mixed cell quantity during training consistently at 500. Through this experimental design, we can observe the effect of the number of cells in mixed samples of training on the deconvolution performance. This will help us understand the performance of scaden and TAPE under different cell mixing scenarios and their applicability with various parameter settings. From Figure 5, it can be observed that when the cell number for generating the training samples is less than the number of test samples, both scaden and TAPE exhibit a noticeable decrease in deconvolution accuracy and stability. This effect is particularly evident when the cell number in the training samples is set to 100. However, when the number of cells in the training samples exceeds 1000, the overall performance of scaden and TAPE tends to stabilize. This phenomenon is likely attributed to the heterogeneity of single-cell data. When the number of cells used to generate the training data is small, the model struggles to learn a stable gene expression profile for deconvolution, resulting in a decrease in effectiveness. In general, bulk data generated in scientific or clinical fields often comprise a vast number of cells, typically far exceeding the order of thousands or tens of thousands. Therefore, we recommend collecting a large amount of high-quality single-cell data and increasing the number of cells when generating training samples when utilizing deep learning methods for deconvolution.

**Figure 5.**
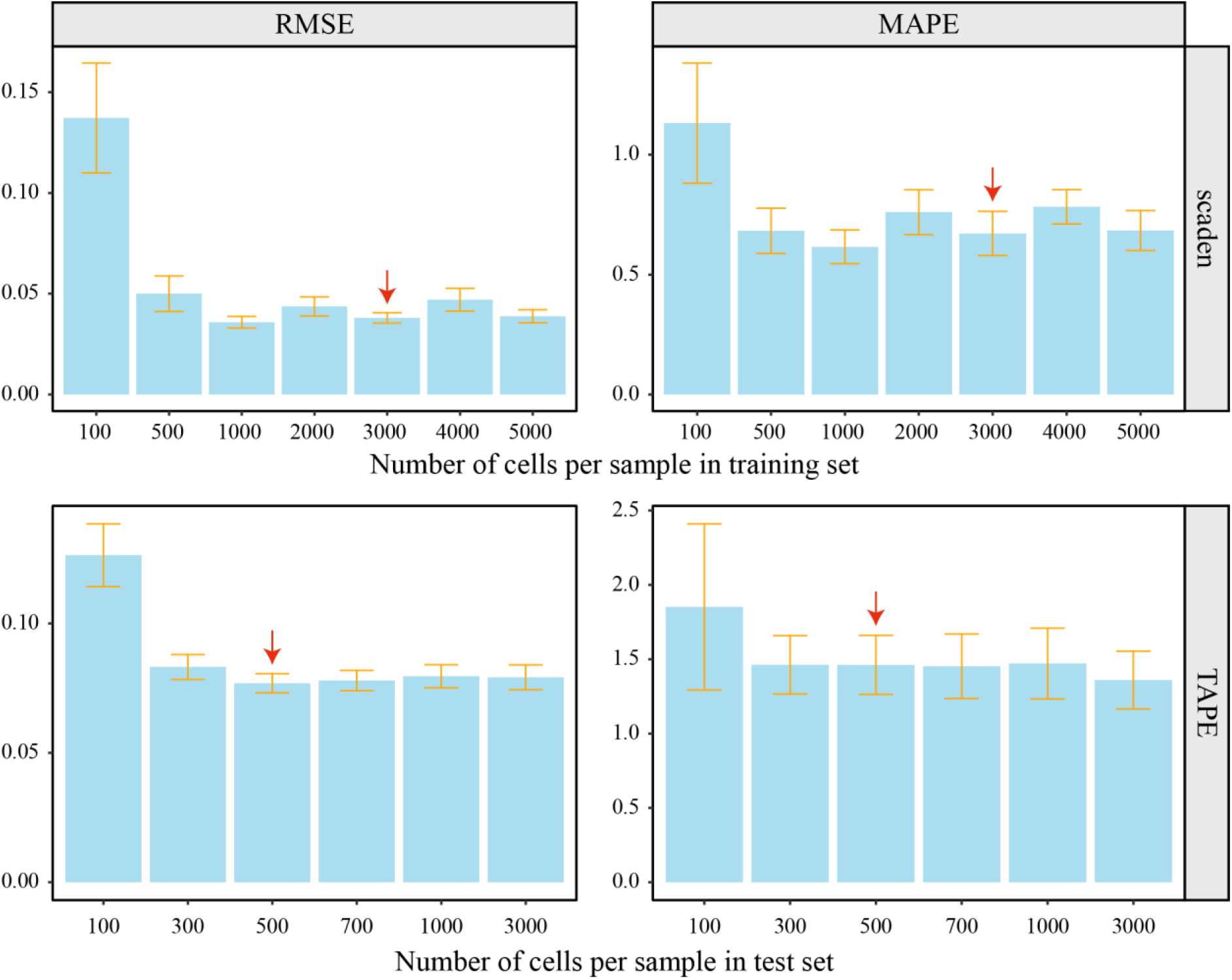
Impact of simulation cell numbers for deconvolution performance. The top panel is the results for scaden and the bottom panel is the results for TAPE. The red arrow means that the number of cells used for generating training and test samples are consistent.

### The potential applications and challenges of cell type deconvolution methods for gene expression data

Cell type deconvolution holds significant promise for applications in clinical medicine. By deciphering the changes in cell proportions, we can gain a deeper understanding of disease progression and pathophysiological mechanisms, providing valuable insights for disease diagnosis, treatment, and prognosis [27,39,53].

Cell type deconvolution can aid in identifying specific biomarkers for particular diseases, enabling early diagnosis of diseases. By analyzing the changes in the proportions of different cell types, disease-related features can be distinguished [38,72]. For instance, an increased proportion of tumor cells in cancer or alterations in the proportions of specific immune cell subsets associated with immune-related disorders [73,74]. In rheumatoid arthritis, variations in the proportions of synovial fibroblasts and immune cell subtypes such as T cells, B cells, and NK cells exist among fibroid, lymphoid, and myeloid types [75]. This provides a crucial foundation for precision medicine, facilitating the development of individualized clinical treatments and interventions.

Cell type deconvolution can also reveal features and patterns of disease progression [20,72]. In many diseases, such as chronic conditions and autoimmune disorders, the changes in the proportions of specific cell types are closely associated with disease progression and severity [4,75–77]. By deciphering the proportion changes of different cell types, we can observe the enrichment or depletion of disease-relevant cell subpopulations, providing insights into disease stages, pathological characteristics, and potential molecular mechanisms. This monitoring can assist in evaluating treatment efficacy and making timely adjustments to treatment plans, maximizing patient survival rates and improving life quality. For example, the alterations in proximal tubule (PT) cells and renal interstitial fibroblasts can reflect the progression of chronic kidney disease [76,77]. Changes in tumor-infiltrating lymphocytes proportions can reflect the clinical efficacy of cancer immunotherapy [78–80]. What’s more, cell type deconvolution provides an important tool for gaining in-depth insights into the pathogenesis of diseases and evaluating drug efficacy during the drug development process [81]. By comparing the changes in cell composition across different disease states, we can uncover the underlying mechanisms, key signaling pathways, and regulatory networks involved in the disease. This aids in identifying novel therapeutic targets and developing innovative treatment strategies, driving advancements in disease research.

Despite the considerable attention and development of deconvolution models, inconsistencies in the deconvolution results still exist among different methods in practical applications. To address this issue, we employed the Unilateral Ureteric Obstruction (UUO) mouse model [33], as used in previous studies [4], to evaluate the 14 deconvolution models mentioned above. The UUO mouse model serves as an experimental simulation of chronic kidney disease (CKD) and consists of three groups of bulk RNA-seq data from renal tissues collected at different time points: sham-operated, 2-day post-ligation, and 8-day post-ligation [33]. In the experiments, various phenomena-related changes in cell type proportions are already known [4,43,82–84]. For example, low-grade inflammation reflects an increase in immune cell proportions, while the development of glomerulosclerosis and renal interstitial fibrosis is associated with a decline in proximal tubule (PT) cell proportion and an increase in fibroblast proportion. As our former work [4], we predicted the proportions of immune cell (summation of macrophage, T lymphocytes, B lymphocytes and nature killer cell), PT cell as well as fibroblast at different stages for each method. According to the results shown in Figure 6, most methods are capable of predicting the trends in cell proportion changes mentioned above. However, there are significant variations in the identified cell proportions. For instance, methods such as scaden, TAPE, FARDEEP, CIBERSORTx, and ARIC accurately capture the changing trends in cell proportions, while CIBERSORT, MOMF, and EPIC exhibit some prediction biases (Figure 6 and Supplementary Figure S10). Nevertheless, several methods demonstrate consistent predictions, including ARIC, FARDEEP, and scaden (Figure 6). Considering the lack of prior knowledge about specific cell type proportions in most applications and the individual differences, it is crucial to employ different deconvolution methods for the same sample. Obtaining consistent results across multiple deconvolution methods will enhance the reliability of the analysis.

**Figure 6.**
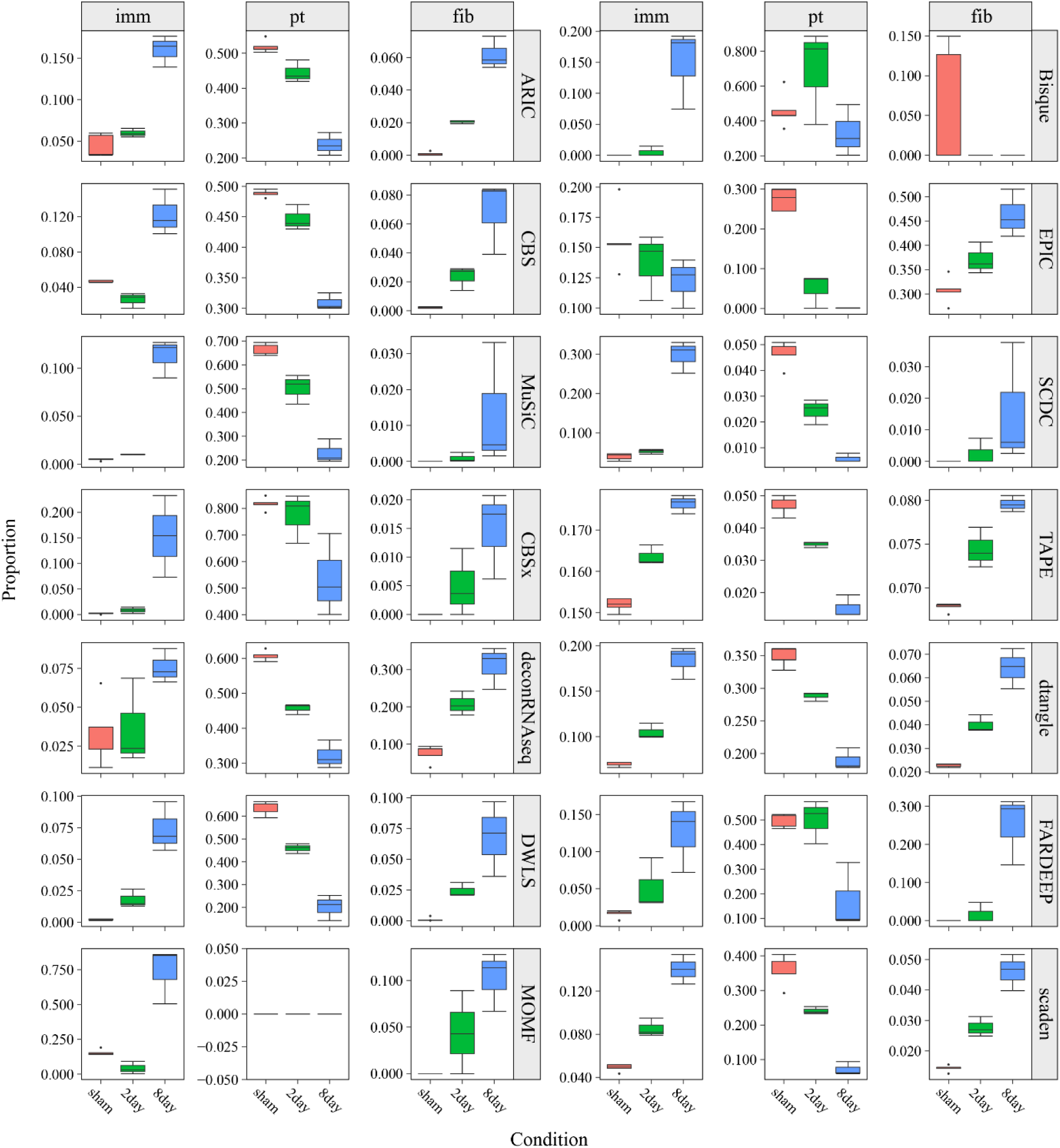
Deconvolution results of each method in the UUO model.

## Discussion

In this study, we developed a unified expression data deconvolution assessment toolkit called Deconer and systematically evaluated 14 reference-based gene expression deconvolution algorithms. Through analysis of both simulated and real-world datasets, it was observed that algorithms such as MuSiC, FARDEEP, DWLS, and ARIC, which incorporate feature selection functions, demonstrated relatively higher accuracy and sensitivity, especially for rare components. Moreover, these methods exhibited consistency in the deconvolution results of real-world data. On the other hand, recent advancements in deep learning techniques, such as scaden and TAPE, have shown enhanced stability and remarkable performance.

It is worth mentioning that cell type deconvolution methods for gene expression data based on deep learning techniques have gained increasing popularity. Leveraging the powerful feature selection and fitting abilities of deep learning models, methods like scaden and TAPE exhibit strong capabilities in addressing batch effect issues[44,45]. However, ensuring high-quality and diverse training data, incorporating prior knowledge of cell type proportions, and optimizing the model with tailored weights for different cell types are critical steps to enhance the stability, accuracy, and effectiveness of these methods. Firstly, it is preferable to have high-quality external single-cell references that ensure an adequate number of cells for each cell type. If the training data used contains high variance and low cell numbers for certain cell types, it can lead to unstable results. Therefore, the quality and quantity of single-cell reference data are crucial for obtaining a stable and accurate deconvolution model. Secondly, when generating training data, it is beneficial to have some prior knowledge of the cell type proportions, especially when dealing with a large number of cell types that need to be identified.

Fully relying on random simulations may not guarantee that the model learns the correct parameter space, particularly for rare components. Lastly, in terms of model improvement, the adoption of different weights for different cell types, as demonstrated by methods like DWLS, has proven to be effective. If weights for the loss function can be adjusted specifically for different cell types within deep learning methods, it will further enhance the effectiveness of fine-tuning the deep learning models.

In addition, the significant discrepancies in the prediction results of different algorithms pose challenges to the practical application of deconvolution algorithms. In real-world applications, the proportions of each cell type are typically unknown, and large variations in the results obtained from different algorithms can reduce the credibility of deconvolution algorithms. In the analysis of cell proportion deconvolution, the differences among different methods may arise from variations in algorithm principles, model assumptions, and data processing. These factors can lead to certain biases in the estimation of proportions for specific cell types. Therefore, it is necessary to consider the results of cell proportion deconvolution in a comprehensive manner, such as through the adoption of ensemble learning with multiple models, and combining with prior knowledge and experimental validation for interpretation and inference. For specific diseases or research proposals, it is advisable to select multiple deconvolution methods for comparison and validation. Although there may be differences among deconvolution methods in predicting the trends of cell proportions, obtaining consistent results through the use of multiple methods will enhance the confidence in cell proportion changes and provide more reliable and comprehensive analytical outcomes.

According to the evaluation analysis in this study and considering previous relevant researches [18,23,24], we provide the following recommendations for reference-based gene expression deconvolution analysis. (1) Ensure that the input data is in the correct data space. Some methods require input in TPM (Transcripts Per Million) format, such as EPIC and ARIC, while others accept raw counts, such as scaden and TAPE. (2) It is preferable for the deconvolution methods to have their own independent feature selection strategies. In our analysis, we found that methods with built-in feature selection strategies, such as FARDEEP and DWLS, consistently performed well. If a method does not have built-in feature selection, we recommend applying strict differential gene selection methods such as DESeq2 or DEsingle before performing deconvolution [85,86]. This not only avoids instability in deconvolution caused by issues such as collinearity but also reduces computational complexity. (3) Whenever possible, use single-cell data as an external reference for deconvolution. High-quality single-cell data plays a crucial role in studying specific cell subpopulations and obtaining more refined deconvolution results. (4) Deconvolution methods should be tailored to the application context. Employing different deconvolution strategies in different scenarios is key to achieving accurate results. For example, for rare components, DWLS or ARIC can be used, while deep learning methods like scaden or TAPE are suitable for addressing batch effect issues. (5) Whenever possible, use multiple deconvolution methods to increase the reliability of the results.

Finally, with the increasing availability of scRNA-seq datasets and the advancements in deep learning, cell proportion deconvolution will become more precise. There is a need for more attention on specific cell types and the development of cell proportion deconvolution algorithms that integrate phenotypic information. The evaluation results presented in this study will enhance the development of deconvolution algorithms, and cell proportion deconvolution will be widely applied in personalized medicine and precision healthcare in the near future.

## Authors’ contributions

The original concept came from XWW, ZPL and WZ. WZ and XLZ conducted the analysis and developed Deconer. QL, LW, XQ, and RG provided guidance for the bioinformatic analysis. All authors participated in interpreting the results. WZ drafted the initial work. All authors contributed to the final draft and critical revisions. All authors reviewed and approved the final manuscript.

## Data Availability

Deconer has been implemented as an open-source R package available at the GitHub repository: https://honchkrow.github.io/Deconer/.

All the public available datasets used in this article are listed in Supplementary Section 1 and accession numbers are listed in Supplementary Table S2. We also provide the processed datasets at: https://honchkrow.github.io/Deconer_dataset/.

## References

[1] Li X, Wang C-Y. From bulk, single-cell to spatial RNA sequencing. Int J Oral Sci 2021;13:1–6.

[2] Chen G, Liu Z-P. Graph attention network for link prediction of gene regulations from single-cell RNA-sequencing data. Bioinformatics 2022:btac559.

[3] Hunt GJ, Freytag S, Bahlo M, Gagnon-Bartsch JA. dtangle: accurate and robust cell type deconvolution. Bioinformatics 2019;35:2093–9.

[4] Zhang W, Xu H, Qiao R, Zhong B, Zhang X, Gu J, et al. ARIC: accurate and robust inference of cell type proportions from bulk gene expression or DNA methylation data. Brief Bioinform 2022;23:bbab362.

[5] Regev A, Teichmann SA, Lander ES, Amit I, Benoist C, Birney E, et al. The Human Cell Atlas. eLife 2017;6:e27041.

[6] Bindea G, Mlecnik B, Tosolini M, Kirilovsky A, Waldner M, Obenauf AC, et al. Spatiotemporal Dynamics of Intratumoral Immune Cells Reveal the Immune Landscape in Human Cancer. Immunity 2013;39:782–95.

[7] Li J, Wei L, Zhang X, Zhang W, Wang H, Zhong B, et al. DISMIR: D eep learning-based noninvasive cancer detection by i ntegrating DNA s equence and methylation information of individual cell-free DNA r eads. Brief Bioinform 2021;22:bbab250.

[8] Galon J, Costes A, Sanchez-Cabo F, Kirilovsky A, Mlecnik B, Lagorce-Pagès C, et al. Type, density, and location of immune cells within human colorectal tumors predict clinical outcome. Science 2006;313:1960–4.

[9] Herzenberg LA, Parks D, Sahaf B, Perez O, Roederer M, Herzenberg LA. The History and Future of the Fluorescence Activated Cell Sorter and Flow Cytometry: A View from Stanford. Clin Chem 2002;48:1819–27.

[10] Newman AM, Steen CB, Liu CL, Gentles AJ, Chaudhuri AA, Scherer F, et al. Determining cell type abundance and expression from bulk tissues with digital cytometry. Nat Biotechnol 2019;37:773–82.

[11] Black CB, Duensing TD, Trinkle LS, Dunlay RT. Cell-Based Screening Using High-Throughput Flow Cytometry. Assay Drug Dev Technol 2011;9:13–20.

[12] Song J, Kuan P-F. A systematic assessment of cell type deconvolution algorithms for DNA methylation data. Brief Bioinform 2022;23:bbac449.

[13] Stoeckius M, Hafemeister C, Stephenson W, Houck-Loomis B, Chattopadhyay PK, Swerdlow H, et al. Large-scale simultaneous measurement of epitopes and transcriptomes in single cells. Nat Methods 2017;14:865–8.

[14] Maesner CC, Almada AE, Wagers AJ. Established cell surface markers efficiently isolate highly overlapping populations of skeletal muscle satellite cells by fluorescence-activated cell sorting. Skelet Muscle 2016;6:35.

[15] Jeong Y, de Andrade e Sousa LB, Thalmeier D, Toth R, Ganslmeier M, Breuer K, et al. Systematic evaluation of cell-type deconvolution pipelines for sequencing-based bulk DNA methylomes. Brief Bioinform 2022;23:bbac248.

[16] Salas LA, Koestler DC, Butler RA, Hansen HM, Wiencke JK, Kelsey KT, et al. An optimized library for reference-based deconvolution of whole-blood biospecimens assayed using the Illumina HumanMethylationEPIC BeadArray. Genome Biol 2018;19:64.

[17] Li H, Sharma A, Luo K, Qin ZS, Sun X, Liu H. DeconPeaker, a Deconvolution Model to Identify Cell Types Based on Chromatin Accessibility in ATAC-Seq Data of Mixture Samples. Front Genet 2020;11.

[18] Jin H, Liu Z. A benchmark for RNA-seq deconvolution analysis under dynamic testing environments. Genome Biol 2021;22:102.

[19] Sun K, Jiang P, Chan KCA, Wong J, Cheng YKY, Liang RHS, et al. Plasma DNA tissue mapping by genome-wide methylation sequencing for noninvasive prenatal, cancer, and transplantation assessments. Proc Natl Acad Sci 2015;112:E5503–12.

[20] Li S, Zeng W, Ni X, Liu Q, Li W, Stackpole ML, et al. Comprehensive tissue deconvolution of cell-free DNA by deep learning for disease diagnosis and monitoring. Proc Natl Acad Sci 2023;120:e2305236120.

[21] Houseman EA, Kile ML, Christiani DC, Ince TA, Kelsey KT, Marsit CJ. Reference-free deconvolution of DNA methylation data and mediation by cell composition effects. BMC Bioinformatics 2016;17:259.

[22] Tang D, Park S, Zhao H. NITUMID: Nonnegative matrix factorization-based Immune-TUmor MIcroenvironment Deconvolution. Bioinformatics 2020;36:1344–50.

[23] Avila Cobos F, Alquicira-Hernandez J, Powell JE, Mestdagh P, De Preter K. Benchmarking of cell type deconvolution pipelines for transcriptomics data. Nat Commun 2020;11:5650.

[24] Mohammadi S, Zuckerman N, Goldsmith A, Grama A. A Critical Survey of Deconvolution Methods for Separating Cell Types in Complex Tissues. Proc IEEE 2017;105:340–66.

[25] Kuhn A, Thu D, Waldvogel HJ, Faull RLM, Luthi-Carter R. Population-specific expression analysis (PSEA) reveals molecular changes in diseased brain. Nat Methods 2011;8:945–7.

[26] Liu R, Holik AZ, Su S, Jansz N, Chen K, Leong HS, et al. Why weight? Modelling sample and observational level variability improves power in RNA-seq analyses. Nucleic Acids Res 2015;43:e97–e97.

[27] Becht E, Giraldo NA, Lacroix L, Buttard B, Elarouci N, Petitprez F, et al. Estimating the population abundance of tissue-infiltrating immune and stromal cell populations using gene expression. Genome Biol 2016;17:218.

[28] Gong T, Hartmann N, Kohane IS, Brinkmann V, Staedtler F, Letzkus M, et al. Optimal Deconvolution of Transcriptional Profiling Data Using Quadratic Programming with Application to Complex Clinical Blood Samples. PLOS ONE 2011;6:e27156.

[29] Linsley PS, Speake C, Whalen E, Chaussabel D. Copy Number Loss of the Interferon Gene Cluster in Melanomas Is Linked to Reduced T Cell Infiltrate and Poor Patient Prognosis. PLoS ONE 2014;9:e109760.

[30] Parsons J, Munro S, Pine PS, McDaniel J, Mehaffey M, Salit M. Using mixtures of biological samples as process controls for RNA-sequencing experiments. BMC Genomics 2015;16:708.

[31] Shen-Orr SS, Tibshirani R, Khatri P, Bodian DL, Staedtler F, Perry NM, et al. Cell type–specific gene expression differences in complex tissues. Nat Methods 2010;7:287–9.

[32] MAQC Consortium, Shi L, Shi L, Reid LH, Jones WD, Shippy R, et al. The MicroArray Quality Control (MAQC) project shows inter- and intraplatform reproducibility of gene expression measurements. Nat Biotechnol 2006;24:1151–61.

[33] Arvaniti E, Moulos P, Vakrakou A, Chatziantoniou C, Chadjichristos C, Kavvadas P, et al. Whole-transcriptome analysis of UUO mouse model of renal fibrosis reveals new molecular players in kidney diseases. Sci Rep 2016;6:26235.

[34] Craciun FL, Bijol V, Ajay AK, Rao P, Kumar RK, Hutchinson J, et al. RNA Sequencing Identifies Novel Translational Biomarkers of Kidney Fibrosis. J Am Soc Nephrol 2016;27:1702.

[35] Vasaikar SV, Straub P, Wang J, Zhang B. LinkedOmics: analyzing multi-omics data within and across 32 cancer types. Nucleic Acids Res 2018;46:D956–63.

[36] Gong T, Szustakowski JD. DeconRNASeq: a statistical framework for deconvolution of heterogeneous tissue samples based on mRNA-Seq data. Bioinformatics 2013;29:1083–5.

[37] Racle J, de Jonge K, Baumgaertner P, Speiser DE, Gfeller D. Simultaneous enumeration of cancer and immune cell types from bulk tumor gene expression data. eLife 2017;6:e26476.

[38] Hao Y, Yan M, Heath BR, Lei YL, Xie Y. Fast and robust deconvolution of tumor infiltrating lymphocyte from expression profiles using least trimmed squares. PLOS Comput Biol 2019;15:e1006976.

[39] Newman AM, Liu CL, Green MR, Gentles AJ, Feng W, Xu Y, et al. Robust enumeration of cell subsets from tissue expression profiles. Nat Methods 2015;12:453–7.

[40] Jew B, Alvarez M, Rahmani E, Miao Z, Ko A, Garske KM, et al. Accurate estimation of cell composition in bulk expression through robust integration of single-cell information. Nat Commun 2020;11:1971.

[41] Tsoucas D, Dong R, Chen H, Zhu Q, Guo G, Yuan G-C. Accurate estimation of cell-type composition from gene expression data. Nat Commun 2019;10:2975.

[42] Sun X, Sun S, Yang S. An Efficient and Flexible Method for Deconvoluting Bulk RNA-Seq Data with Single-Cell RNA-Seq Data. Cells 2019;8:1161.

[43] Wang X, Park J, Susztak K, Zhang NR, Li M. Bulk tissue cell type deconvolution with multi-subject single-cell expression reference. Nat Commun 2019;10:380.

[44] Menden K, Marouf M, Oller S, Dalmia A, Magruder DS, Kloiber K, et al. Deep learning–based cell composition analysis from tissue expression profiles. Sci Adv 2020;6:eaba2619.

[45] Chen Y, Wang Y, Chen Y, Cheng Y, Wei Y, Li Y, et al. Deep autoencoder for interpretable tissue-adaptive deconvolution and cell-type-specific gene analysis. Nat Commun 2022;13:6735.

[46] Dong M, Thennavan A, Urrutia E, Li Y, Perou CM, Zou F, et al. SCDC: bulk gene expression deconvolution by multiple single-cell RNA sequencing references. Brief Bioinform 2021;22:416–27.

[47] Han X, Wang R, Zhou Y, Fei L, Sun H, Lai S, et al. Mapping the Mouse Cell Atlas by Microwell-Seq. Cell 2018;172:1091–1107.e17.

[48] Bredikhin D, Kats I, Stegle O. MUON: multimodal omics analysis framework. Genome Biol 2022;23:42.

[49] Law CW, Chen Y, Shi W, Smyth GK. voom: precision weights unlock linear model analysis tools for RNA-seq read counts. Genome Biol 2014;15:R29.

[50] Hu T, Wei L, Li S, Cheng T, Zhang X, Wang X. Single-cell Transcriptomes Reveal Characteristics of MicroRNAs in Gene Expression Noise Reduction. Genomics Proteomics Bioinformatics 2021;19:394–407.

[51] Yu X, Chen YA, Conejo-Garcia JR, Chung CH, Wang X. Estimation of immune cell content in tumor using single-cell RNA-seq reference data. BMC Cancer 2019;19:715.

[52] Pei G, Wang Y-Y, Simon LM, Dai Y, Zhao Z, Jia P. Gene expression imputation and cell-type deconvolution in human brain with spatiotemporal precision and its implications for brain-related disorders. Genome Res 2021;31:146–58.

[53] Chu T, Wang Z, Pe’er D, Danko CG. Cell type and gene expression deconvolution with BayesPrism enables Bayesian integrative analysis across bulk and single-cell RNA sequencing in oncology. Nat Cancer 2022;3:505–17.

[54] Pournara AV, Miao Z, Beker O, Brazma A, Papatheodorou I. Power analysis of cell-type deconvolution methods across tissues. In Review; 2023.

[55] Li Y, Luo P, Lu Y, Wu F-X. Identifying cell types from single-cell data based on similarities and dissimilarities between cells. BMC Bioinformatics 2021;22:255.

[56] Watanabe K, Umićević Mirkov M, de Leeuw CA, van den Heuvel MP, Posthuma D. Genetic mapping of cell type specificity for complex traits. Nat Commun 2019;10:3222.

[57] Gao C, Sun H, Wang T, Tang M, Bohnen NI, Müller MLTM, et al. Model-based and Model-free Machine Learning Techniques for Diagnostic Prediction and Classification of Clinical Outcomes in Parkinson’s Disease. Sci Rep 2018;8:7129.

[58] Liu Y-T, Sun Z-J. Turning cold tumors into hot tumors by improving T-cell infiltration. Theranostics 2021;11:5365–86.

[59] Chakravarthy A, Furness A, Joshi K, Ghorani E, Ford K, Ward MJ, et al. Pan-cancer deconvolution of tumour composition using DNA methylation. Nat Commun 2018;9:3220.

[60] Grabovska Y, Mackay A, O’Hare P, Crosier S, Finetti M, Schwalbe EC, et al. Pediatric pan-central nervous system tumor analysis of immune-cell infiltration identifies correlates of antitumor immunity. Nat Commun 2020;11:4324.

[61] Gentles AJ, Newman AM, Liu CL, Bratman SV, Feng W, Kim D, et al. The prognostic landscape of genes and infiltrating immune cells across human cancers. Nat Med 2015;21:938–45.

[62] Teixeira L, Rothé F, Ignatiadis M, Sotiriou C. Breast Cancer Immunology. Oncol Times 2016;38:18.

[63] Moufarrej MN, Vorperian SK, Wong RJ, Campos AA, Quaintance CC, Sit RV, et al. Early prediction of preeclampsia in pregnancy with cell-free RNA. Nature 2022;602:689–94.

[64] Vorperian SK, Moufarrej MN, Quake SR. Cell types of origin of the cell-free transcriptome. Nat Biotechnol 2022;40:855–61.

[65] Tosevska A, Morselli M, Basak SK, Avila L, Mehta P, Wang MB, et al. Cell-Free RNA as a Novel Biomarker for Response to Therapy in Head & Neck Cancer. Front Oncol 2022;12.

[66] Moufarrej MN, Bianchi DW, Shaw GM, Stevenson DK, Quake SR. Noninvasive Prenatal Testing Using Circulating DNA and RNA: Advances, Challenges, and Possibilities. Annu Rev Biomed Data Sci 2023;6:null.

[67] Larson MH, Pan W, Kim HJ, Mauntz RE, Stuart SM, Pimentel M, et al. A comprehensive characterization of the cell-free transcriptome reveals tissue- and subtype-specific biomarkers for cancer detection. Nat Commun 2021;12:2357.

[68] Tran KA, Kondrashova O, Bradley A, Williams ED, Pearson JV, Waddell N. Deep learning in cancer diagnosis, prognosis and treatment selection. Genome Med 2021;13:152.

[69] Bassiouni R, Gibbs LD, Craig DW, Carpten JD, McEachron TA. Applicability of spatial transcriptional profiling to cancer research. Mol Cell 2021;81:1631–9.

[70] Adlung L, Cohen Y, Mor U, Elinav E. Machine learning in clinical decision making. Med 2021;2:642–65.

[71] Goldman SL, MacKay M, Afshinnekoo E, Melnick AM, Wu S, Mason CE. The Impact of Heterogeneity on Single-Cell Sequencing. Front Genet 2019;10.

[72] Zhang W, Wei L, Huang J, Zhong B, Li J, Xu H, et al. cfDNApipe: A comprehensive quality control and analysis pipeline for cell-free DNA high-throughput sequencing data. Bioinformatics 2021:btab413.

[73] Li T, Fan J, Wang B, Traugh N, Chen Q, Liu JS, et al. TIMER: A Web Server for Comprehensive Analysis of Tumor-Infiltrating Immune Cells. Cancer Res 2017;77:e108–10.

[74] Li T, Fu J, Zeng Z, Cohen D, Li J, Chen Q, et al. TIMER2.0 for analysis of tumor-infiltrating immune cells. Nucleic Acids Res 2020;48:W509–14.

[75] Micheroli R, Elhai M, Edalat S, Frank-Bertoncelj M, Bürki K, Ciurea A, et al. Role of synovial fibroblast subsets across synovial pathotypes in rheumatoid arthritis: a deconvolution analysis. RMD Open 2022;8:e001949.

[76] Kirita Y, Wu H, Uchimura K, Wilson PC, Humphreys BD. Cell profiling of mouse acute kidney injury reveals conserved cellular responses to injury. Proc Natl Acad Sci 2020;117:15874–83.

[77] Park J, Shrestha R, Qiu C, Kondo A, Huang S, Werth M, et al. Single-cell transcriptomics of the mouse kidney reveals potential cellular targets of kidney disease. Science 2018;360:758–63.

[78] Liu CC, Steen CB, Newman AM. Computational approaches for characterizing the tumor immune microenvironment. Immunology 2019;158:70–84.

[79] Zhang Z, Wiencke JK, Kelsey KT, Koestler DC, Christensen BC, Salas LA. HiTIMED: hierarchical tumor immune microenvironment epigenetic deconvolution for accurate cell type resolution in the tumor microenvironment using tumor-type-specific DNA methylation data. J Transl Med 2022;20:516.

[80] Oswald E, Bug D, Grote A, Lashuk K, Bouteldja N, Lenhard D, et al. Immune cell infiltration pattern in non-small cell lung cancer PDX models is a model immanent feature and correlates with a distinct molecular and phenotypic make-up. J Immunother Cancer 2022;10:e004412.

[81] Zhao W, Dovas A, Spinazzi EF, Levitin HM, Banu MA, Upadhyayula P, et al. Deconvolution of cell type-specific drug responses in human tumor tissue with single-cell RNA-seq. Genome Med 2021;13:82.

[82] Chevalier RL. The proximal tubule is the primary target of injury and progression of kidney disease: role of the glomerulotubular junction. Am J Physiol Renal Physiol 2016;311:F145–161.

[83] Neagu M, Zipeto D, Popescu ID. Inflammation in Cancer: Part of the Problem or Part of the Solution? J Immunol Res 2019;2019:5403910.

[84] Akchurin OM, Kaskel F. Update on inflammation in chronic kidney disease. Blood Purif 2015;39:84–92.

[85] Miao Z, Deng K, Wang X, Zhang X. DEsingle for detecting three types of differential expression in single-cell RNA-seq data. Bioinformatics 2018;34:3223–4.

[86] Love MI, Huber W, Anders S. Moderated estimation of fold change and dispersion for RNA-seq data with DESeq2. Genome Biol 2014;15:550.

