## Supplemental Figures for "Deconer: A comprehensive and systematic evaluation toolkit for reference-based cell type deconvolution algorithms using gene expression data"

### 1 Package "Deconer"

#### 1.1 Introduction

Cell type proportion is related to phenotypes or diseases [1,2]. Therefore, quantifying cell or tissue proportions is important for understanding the mechanisms in biological processes. Here, we proposed a cell type deconvolution evaluating toolkit named 'Deconer' to perform comprehensive and systematic evaluation of different algorithms.

Deconer consists of 6 main part functions as below.

- Pseudo bulk data generation (from massive bulk data and single cell data).
- Stability analysis under different types of noise.
- Rare component analysis.
- Unknown component analysis.
- Comprehensive evaluation metrics as well as pretty figure generation.
- Well-characterized datasets for deconvolution utilities.

For more information, please see the following pages.

- Deconer homepage
- Deconer manual
- Deconer vignettes
- Deconer dataset page

#### 1.2 Installation

Deconer is based on R and can be easily installed on Windows, Linux as well as MAC OS.

First, users should install R >= 4.1.0.

Next, install devtools and Deconer.

```
# install devtools
install.packages('devtools')

# install the decone package
devtools::install_github('Honchkrow/decone')

# load decone
library(decone)
```

#### 1.3 Functions used in this study

We generated all the simulated data using Deconer package, including pseudo bulk data generated from massive RNA-seq data as well as from scRNA-seq data (see Deconer homepage Section 3: Gnerating Pseudo Bulk Data). For noise analysis, please see Deconer homepage Section 4: Noise Analysis. For rare component analysis, please see Deconer homepage Section 5: Rare Component Analysis. In addition, most of the figures are also generated directly from Deconer (see Deconer homepage Code Demo 1: Evaluating The Deconvolution Results In A Simple Manner and Code Demo 2: Evaluating The Deconvolution Results For Multiple Method).

#### 1.4 Datasets

Besides, we collected well-characterized deconvolution datasets for users. Some of them are known cell-type proportions. Some datasets without know the true proportion, but the related phenotype can be accessed. We provide the processed datasets with bulk data, reference data as well as true proportions, which means that these datasets can be used directly. We summarized the datasets in the following table S1.

**Table S1**. Summary of the dataset used in Deconer

| NO. | Dataset Name | Data type | Reference Type | Proportion | Source |
|---|---|---|---|---|---|
| 1 | Abbas | Microarray | Bulk | known | [3] |
| 2 | Becht | Microarray | Bulk | known | [4] |
| 3 | Gong | Microarray | Bulk | known | [5] |
| 4 | Kuhn | Microarray | Bulk | known | [6] |
| 5 | Linsley | RNA-seq | Bulk | known | [7] |
| 6 | Liu | RNA-seq | Bulk | known | [8] |
| 7 | Parsons | RNA-seq | Bulk | known | [9] |
| 8 | Shen-Orr | Microarray | Bulk | known | [10] |
| 9 | Shi | Microarray | Bulk | known | [11] |
| 10 | T2D | RNA-seq | Single Cell | unknown | [12] |
| 11 | TCGA_LUSC | RNA-seq | Bulk | unknown | [13] |
| 12 | TCGA_OV | RNA-seq | Bulk | unknown | [13] |
| 13 | kidney_Arvaniti | RNA-seq | Single Cell | unknown | [14] |
| 14 | kidney_Arvaniti_TPM | RNA-seq | Bulk | unknown | [14] |
| 15 | kidney_Craciun | RNA-seq | Single Cell | unknown | [15] |
| 16 | kidney_Craciun_TPM | RNA-seq | Bulk | unknown | [15] |
| 17 | TCGA 35 cancer datasets | RNA-seq | Bulk | unknown | [13] |

All the datasets can be downloaded from Deconer dataset page.

### 2 Datasets Processing

#### 2.1 Bulk RNA-seq

We collected 233 high-quality bulk RNA-seq samples to simulate different stromal populations, including immune populations, in this study. Drawing inspiration from the Tumor Deconvolution DREAM Challenge organized by SYNAPSE, we created two distinct scenarios referred to as "coarse" and "fine". In the coarse scenario, the mixed samples comprised 8 different cell types: B cells, CD4 T cells, CD8 T cells, endothelial cells, macrophages, monocytes, neutrophils, and NK cells. In the fine scenario, the mixed samples encompassed 14 different cell types: memory B cells, naive B cells, memory CD4 T cells, active CD4 T cells, regulatory T cells, memory CD8 T cells, naive CD8 T cells, NK cells, neutrophils, monocytes, myeloid dendritic cells, macrophages, fibroblasts, and endothelial cells. During data collection, we made efforts to ensure that each cell type originated from at least two different studies and that all samples were paired-end data. This approach aimed to mitigate batch effects arising from variations between single- and paired-end sequencing experiments. It is evident that in the "fine" dataset, various cell types are further subdivided into subtypes, which often share similar biological functions. Subtypes with similar functions generally exhibit comparable gene expression profiles, making it more challenging to accurately estimate the proportions of these subtype cells.

For data processing, we employed the following RNA-seq data processing workflow. In simple terms, we downloaded the fastq files for all samples and used Adapterremoval to remove adapter sequences [16]. Then, we utilized Hisat2 to align all transcriptome samples to the hg19 reference genome with the parameters "-q --no-mixed --no-discordant", and obtained bam files using samtools [17,18]. During this process, QuaCRS and iSeqQC were employed for quality control, and low-quality and low-depth data were directly discarded [19,20]. Subsequently, we converted the bam files into gene expression counts using featureCounts [21].

#### 2.2 scRNA-seq

We collected 10K single-cell data of human PBMCs from 10x Genomics. For this dataset, we used muon to annotate 13 reliable cell types [22]. Additionally, we curated a unique single-cell dataset for the generation of mixed samples. This dataset consisted of seven different tissues from fetal mouse [23]. Given the significant expression profile variations across these seven tissues, the samples generated from this dataset served as a fair benchmark for evaluating the capabilities of different algorithms. In addition to the simulated data, we also employed a well-characterized mouse kidney dataset to test all the deconvolution methods discussed in this paper. This dataset represents real data with unknown proportions, and we assessed the deconvolution methods by comparing the inferred cell proportions with known biological phenotypes or processes.

##### 2.3 Data transformation and normalization

During the analysis process, all data were transformed into the required input format for each respective method. For instance, methods like EPIC and ARIC require TPM as input, while dtangle requires log-transformed data [2,24,25]. deconRNAseq, on the other hand, requires data to be maintained in linear space to preserve the assumptions of the linear regression model [26]. Regarding data normalization, scaden and TAPE, being deep learning methods, handle the min_max_scale internally [27,28]. Previous studies have indicated that the zero mean and unit variance data standardization used by CIBERSORT may introduce estimation bias [29]. Therefore, we did not include data normalization beyond the required transformations for each algorithm.

#### 3 Signature Genes Selection

Multiple studies have indicated that differentially expressed genes play a crucial role in cell proportion deconvolution [2,24]. Therefore, we employed DEseq2 and DEsingle for signature gene selection [30,31]. Specifically, DEseq2 was used for differential gene screening in bulk data, while DEsingle was employed for single-cell data. In general, our selection of differentially expressed genes followed the following principles. Firstly, genes with low expression quality were excluded. Secondly, for methods like scaden and TAPE that have their own high variance gene selection strategies, which explicitly state that separate differential gene screening is not required, we included all genes as input [27,28]. Additionally, our differential gene selection was based on the "one_VS_others" strategy, meaning that when selecting differential genes for a particular cell type, all other cell types were treated as the control samples. For DEseq2, we utilized parameter settings with a p-value of 0.001, fold change of 4, and baseMean greater than 10 and less than 1000. As for DEsingle, we extracted differentially expressed genes of the "DEg" type and set the p-value threshold at 0.0001 and fold change at 4. For algorithms that do not provide their own feature selection strategies, we directly employed all identified differentially expressed genes as signature genes for deconvolution.

#### 4 *In Silico* Simulation

For bulk data, we adopted the data partitioning strategy of ARIC [2]. As for scRNA-seq data, we initially divided the data for each cell type into a train set (70%) and a test set (30%) based on the number of cells. To generate pseudo bulk data, we first created random cell proportions and then sampled from the test set based on the predetermined sample cell counts, like 500 or 3000. Since in many cases the single-cell data for specific cell types may not be abundant, Deconer incorporates a parameter that allows for repeated cell sampling.

### 5 Reference


1. Wang X, Park J, Susztak K, et al. Bulk tissue cell type deconvolution with multi-subject single-cell expression reference. Nat. Commun. 2019; 10:380

2. Zhang W, Xu H, Qiao R, et al. ARIC: accurate and robust inference of cell type proportions from bulk gene expression or DNA methylation data. Brief. Bioinform. 2022; 23:bbab362

3. Abbas AR, Wolslegel K, Seshasayee D, et al. Deconvolution of Blood Microarray Data Identifies Cellular Activation Patterns in Systemic Lupus Erythematosus. PLoS ONE 2009; 4:e6098

4. Becht E, Giraldo NA, Lacroix L, et al. Estimating the population abundance of tissue-infiltrating immune and stromal cell populations using gene expression. Genome Biol. 2016; 17:218

5. Gong T, Hartmann N, Kohane IS, et al. Optimal Deconvolution of Transcriptional Profiling Data Using Quadratic Programming with Application to Complex Clinical Blood Samples. PLOS ONE 2011; 6:e27156

6. Kuhn A, Thu D, Waldvogel HJ, et al. Population-specific expression analysis (PSEA) reveals molecular changes in diseased brain. Nat. Methods 2011; 8:945–947

7. Linsley PS, Speake C, Whalen E, et al. Copy Number Loss of the Interferon Gene Cluster in Melanomas Is Linked to Reduced T Cell Infiltrate and Poor Patient Prognosis. PLoS ONE 2014; 9:e109760

8. Liu R, Holik AZ, Su S, et al. Why weight? Modelling sample and observational level variability improves power in RNA-seq analyses. Nucleic Acids Res. 2015; 43:e97–e97

9. Parsons J, Munro S, Pine PS, et al. Using mixtures of biological samples as process controls for RNA-sequencing experiments. BMC Genomics 2015; 16:708

10. Shen-Orr SS, Tibshirani R, Khatri P, et al. Cell type–specific gene expression differences in complex tissues. Nat. Methods 2010; 7:287–289

11. MAQC Consortium, Shi L, Shi L, et al. The MicroArray Quality Control (MAQC) project shows inter- and intraplatform reproducibility of gene expression measurements. Nat. Biotechnol. 2006; 24:1151–1161

12. Fadista J, Vikman P, Laakso EO, et al. Global genomic and transcriptomic analysis of human pancreatic islets reveals novel genes influencing glucose metabolism. Proc. Natl. Acad. Sci. 2014; 111:13924–13929

13. Vasaikar SV, Straub P, Wang J, et al. LinkedOmics: analyzing multi-omics data within and across 32 cancer types. Nucleic Acids Res. 2018; 46:D956–D963

14. Arvaniti E, Moulos P, Vakrakou A, et al. Whole-transcriptome analysis of UUO mouse model of renal fibrosis reveals new molecular players in kidney diseases. Sci. Rep. 2016; 6:26235

15. Craciun FL, Bijol V, Ajay AK, et al. RNA Sequencing Identifies Novel Translational Biomarkers of Kidney Fibrosis. J. Am. Soc. Nephrol. 2016; 27:1702

16. Schubert M, Lindgreen S, Orlando L. AdapterRemoval v2: rapid adapter trimming, identification, and read merging. BMC Res. Notes 2016; 9:88

17. Danecek P, Bonfield JK, Liddle J, et al. Twelve years of SAMtools and BCFtools. GigaScience 2021; 10:giab008

18. Kim D, Paggi JM, Park C, et al. Graph-based genome alignment and genotyping with HISAT2 and HISAT-genotype. Nat. Biotechnol. 2019; 37:907–915

19. Kroll KW, Mokaram NE, Pelletier AR, et al. Quality Control for RNA-Seq (QuaCRS): An Integrated Quality Control Pipeline. Cancer Inform. 2014; 13:7–14

20. Kumar G, Ertel A, Feldman G, et al. iSeqQC: a tool for expression-based quality control in RNA sequencing. BMC Bioinformatics 2020; 21:56

21. Liao Y, Smyth GK, Shi W. featureCounts: an efficient general purpose program for assigning sequence reads to genomic features. Bioinformatics 2014; 30:923–930

22. Bredikhin D, Kats I, Stegle O. MUON: multimodal omics analysis framework. Genome Biol. 2022; 23:42

23. Han X, Wang R, Zhou Y, et al. Mapping the Mouse Cell Atlas by Microwell-Seq. Cell 2018; 172:1091-1107.e17

24. Racle J, de Jonge K, Baumgaertner P, et al. Simultaneous enumeration of cancer and immune cell types from bulk tumor gene expression data. eLife 2017; 6:e26476

25. Hunt GJ, Freytag S, Bahlo M, et al. dtangle: accurate and robust cell type deconvolution. Bioinformatics 2019; 35:2093–2099

26. Gong T, Szustakowski JD. DeconRNASeq: a statistical framework for deconvolution of heterogeneous tissue samples based on mRNA-Seq data. Bioinformatics 2013; 29:1083–1085

27. Menden K, Marouf M, Oller S, et al. Deep learning–based cell composition analysis from tissue expression profiles. Sci. Adv. 2020; 6:eaba2619

28. Chen Y, Wang Y, Chen Y, et al. Deep autoencoder for interpretable tissue-adaptive deconvolution and cell-type-specific gene analysis. Nat. Commun. 2022; 13:6735

29. Hao Y, Yan M, Heath BR, et al. Fast and robust deconvolution of tumor infiltrating lymphocyte from expression profiles using least trimmed squares. PLOS Comput. Biol. 2019; 15:e1006976

30. Love MI, Huber W, Anders S. Moderated estimation of fold change and dispersion for RNA-seq data with DESeq2. Genome Biol. 2014; 15:550

31. Miao Z, Deng K, Wang X, et al. DEsingle for detecting three types of differential expression in single-cell RNA-seq data. Bioinformatics 2018; 34:3223–3224

1

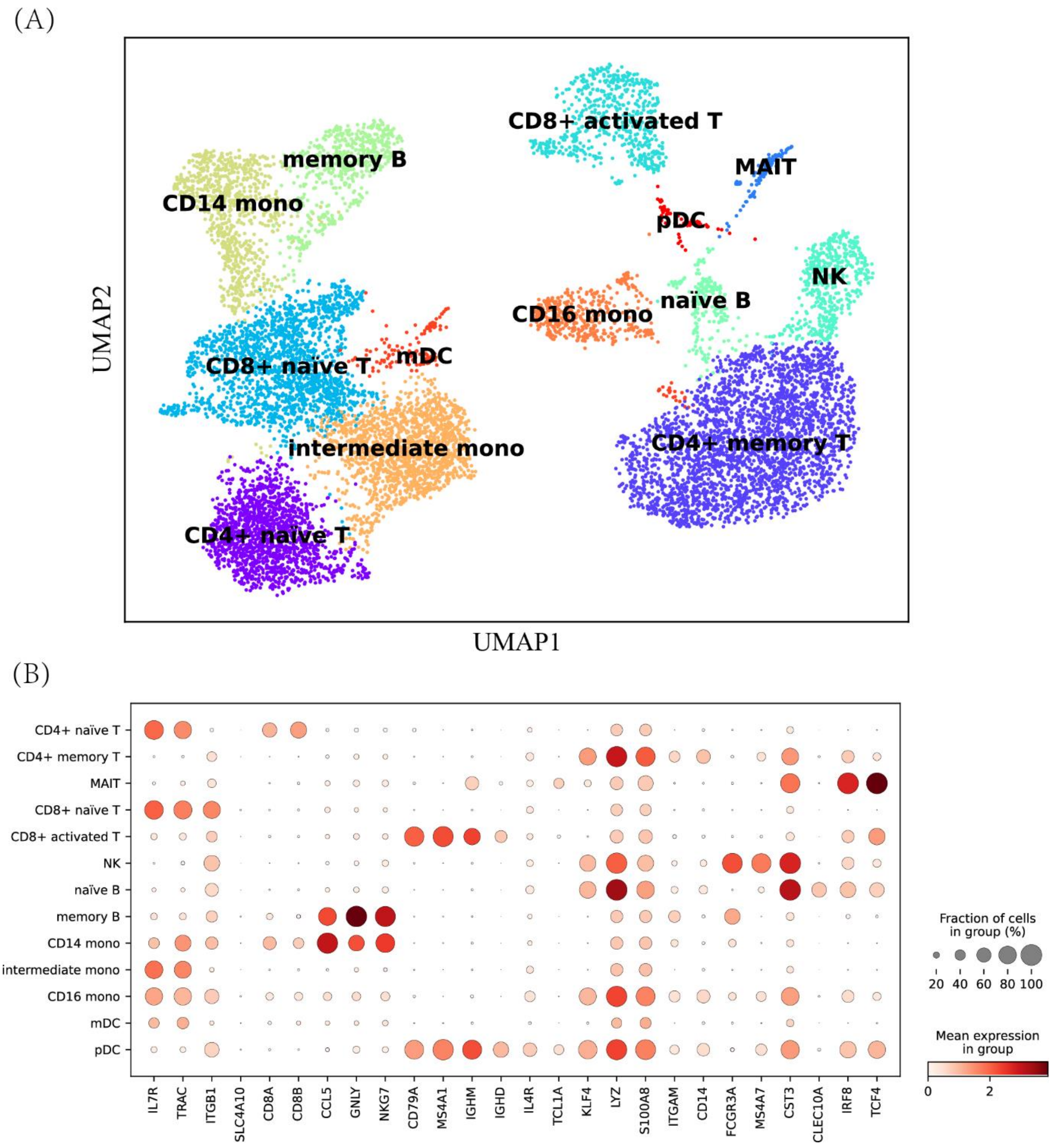


**Figure S1.** Cell annotation for human_PBMC dataset. (A) UMAP plot for cell clusters. (B) The selected marker genes for each cell types.

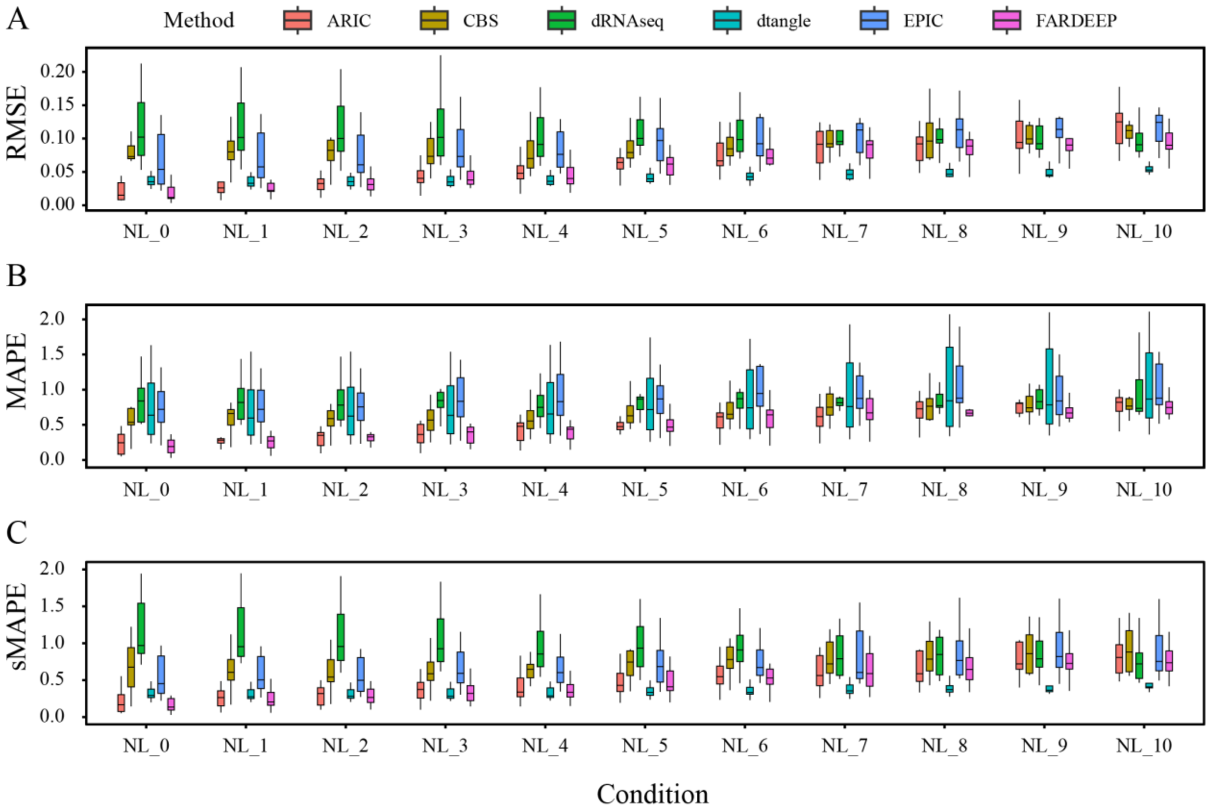


**Figure S2.** Performance metrics of different algorithms on dataset coarse_bulk. RMSE, MAPE as well as sMAPE are illustrated in (A), (B) and (C) respectively. We set $p_t$ in a range from 0.1 to 1 to control noise level (NL_1 to NL_10). NL_0 denotes the absence of noise incorporation. CBS: CIBERSORT, dRNAseq: DeconRNA-seq.

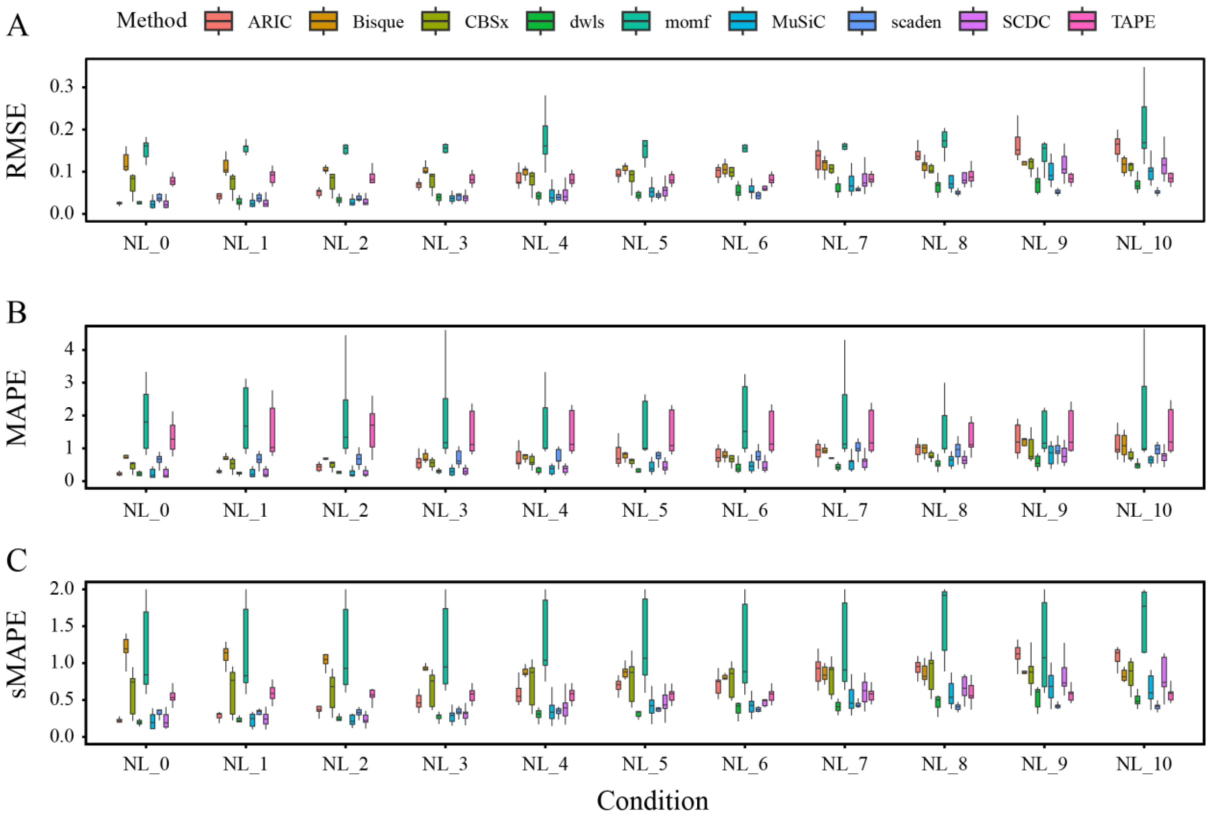


**Figure S3.** Performance metrics of different algorithms on dataset mouse_tissue. RMSE, MAPE as well as sMAPE are illustrated in (A), (B) and (C) respectively. We set $p_t$ in a range from 0.1 to 1 to control noise level (NL_1 to NL_10). NL_0 denotes the absence of noise incorporation. CBSx: CIBERSORTx.

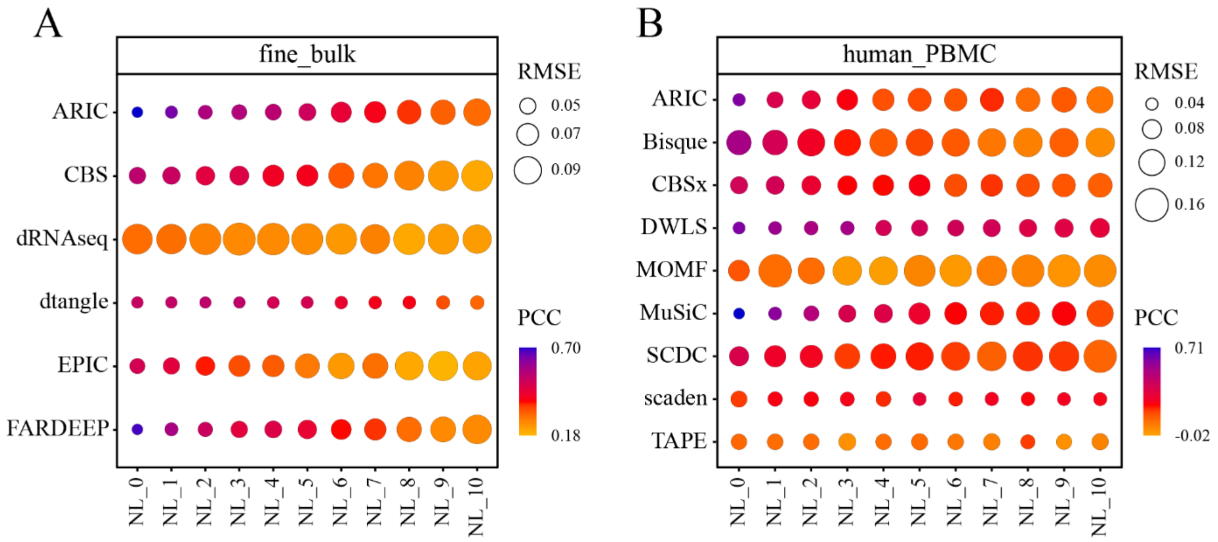


**Figure S4.** Stability testing of various deconvolution methods in scenarios with many cell types. (A) and (B) are the RMSE and PCC for methods employing bulk data and single cell data as a reference respectively. We set $p_t$ in a range from 0.1 to 1 to control noise level (NL_1 to NL_10). NL_0 denotes the absence of noise incorporation. CBS: CIBERSORT, dRNAseq: DeconRNA-seq. CBSx: CIBERSORTx.

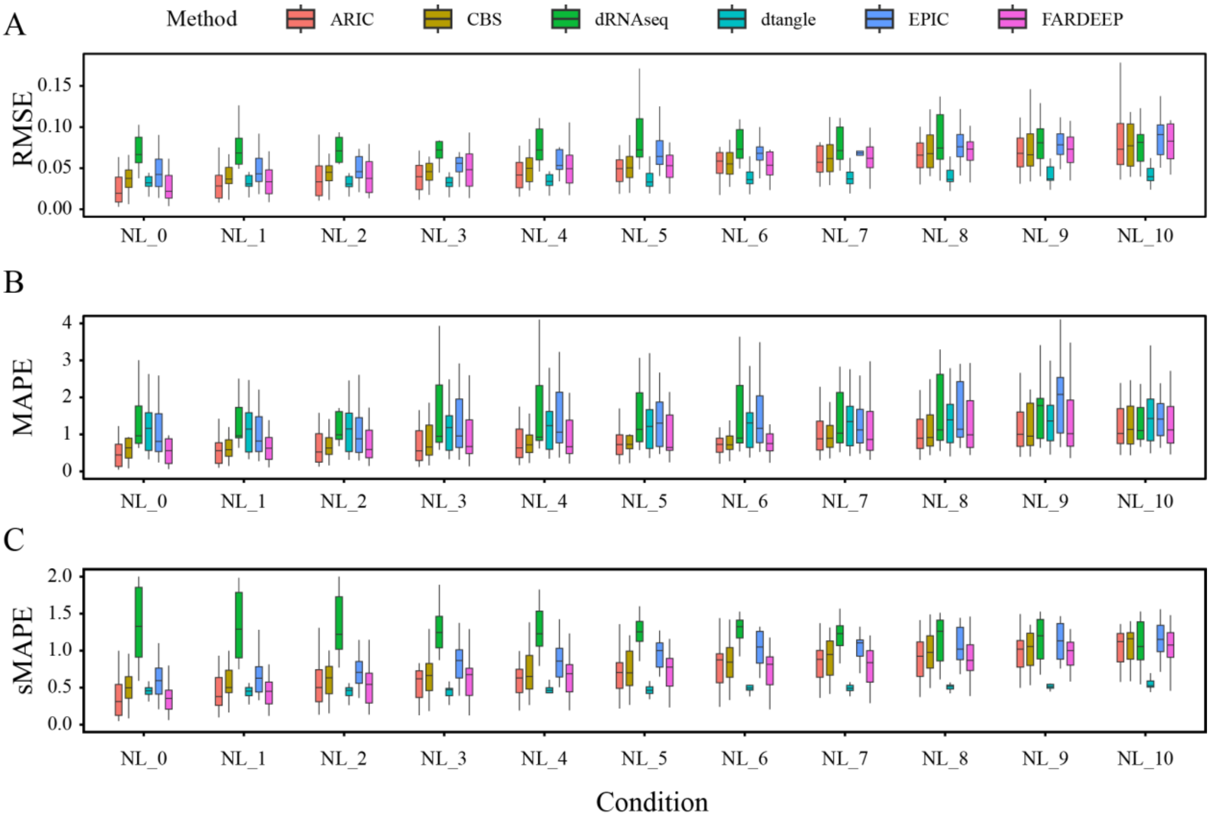


**Figure S5.** Performance metrics of different algorithms on dataset fine_bulk. RMSE, MAPE as well as sMAPE are illustrated in (A), (B) and (C) respectively. We set $p_t$ in a range from 0.1 to 1 to control noise level (NL_1 to NL_10). NL_0 denotes the absence of noise incorporation. CBS: CIBERSORT, dRNAseq: DeconRNA-seq.

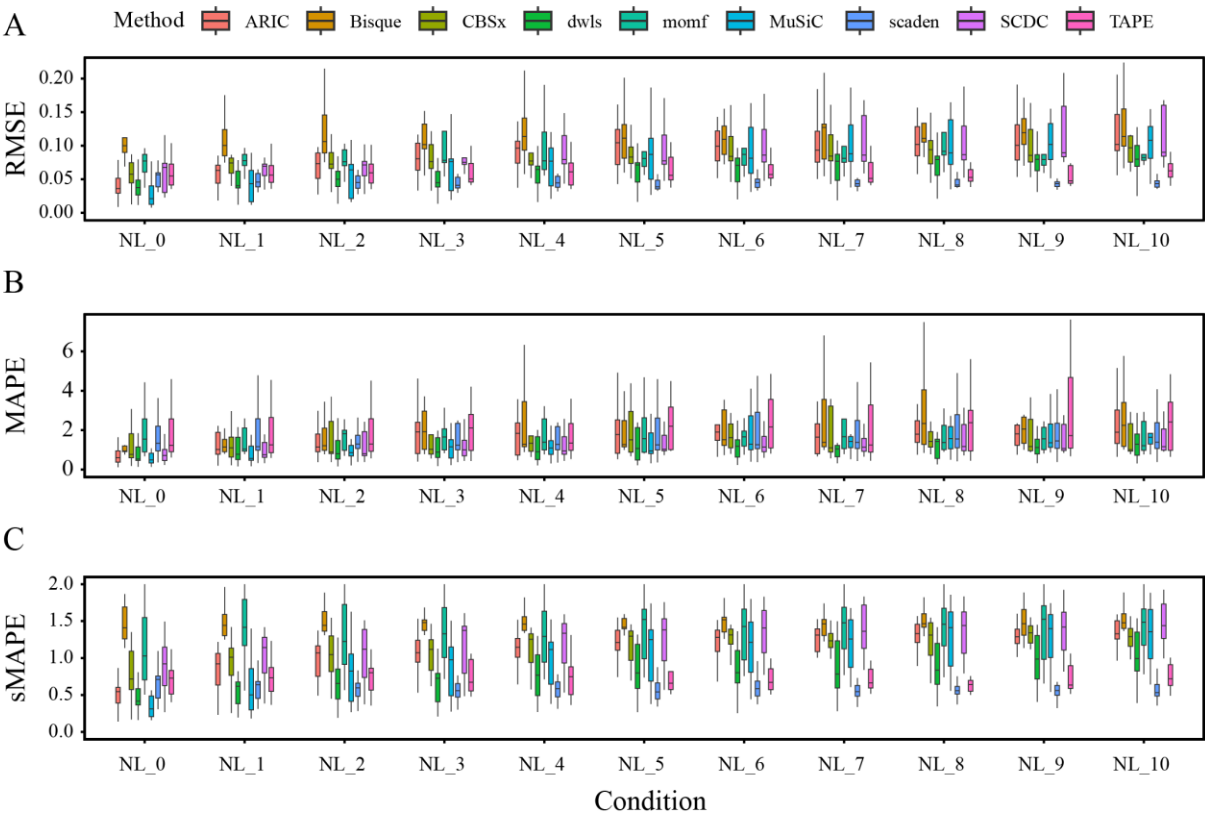


**Figure S6.** Performance metrics of different algorithms on dataset human_PBMC. RMSE, MAPE as well as sMAPE are illustrated in (A), (B) and (C) respectively. We set $p_t$ in a range from 0.1 to 1 to control noise level (NL_1 to NL_10). NL_0 denotes the absence of noise incorporation. CBSx: CIBERSORTx.

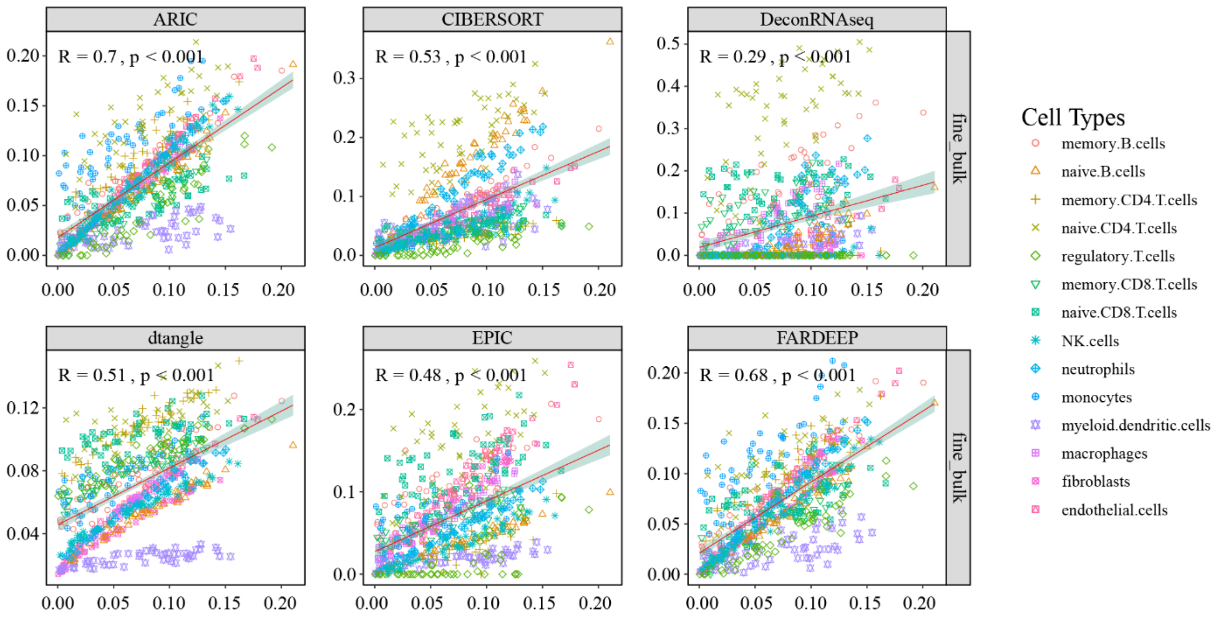


**Figure S7.** Deconvolution results for fine_bulk datasets. This figure corresponds to NL_0 in the figure S5, which means that noise is not added in this test. The datasets are annotated on the right side of the graph, and the names of each method are labelled above their respective graphs. The light green area represents the 95% confidence interval.

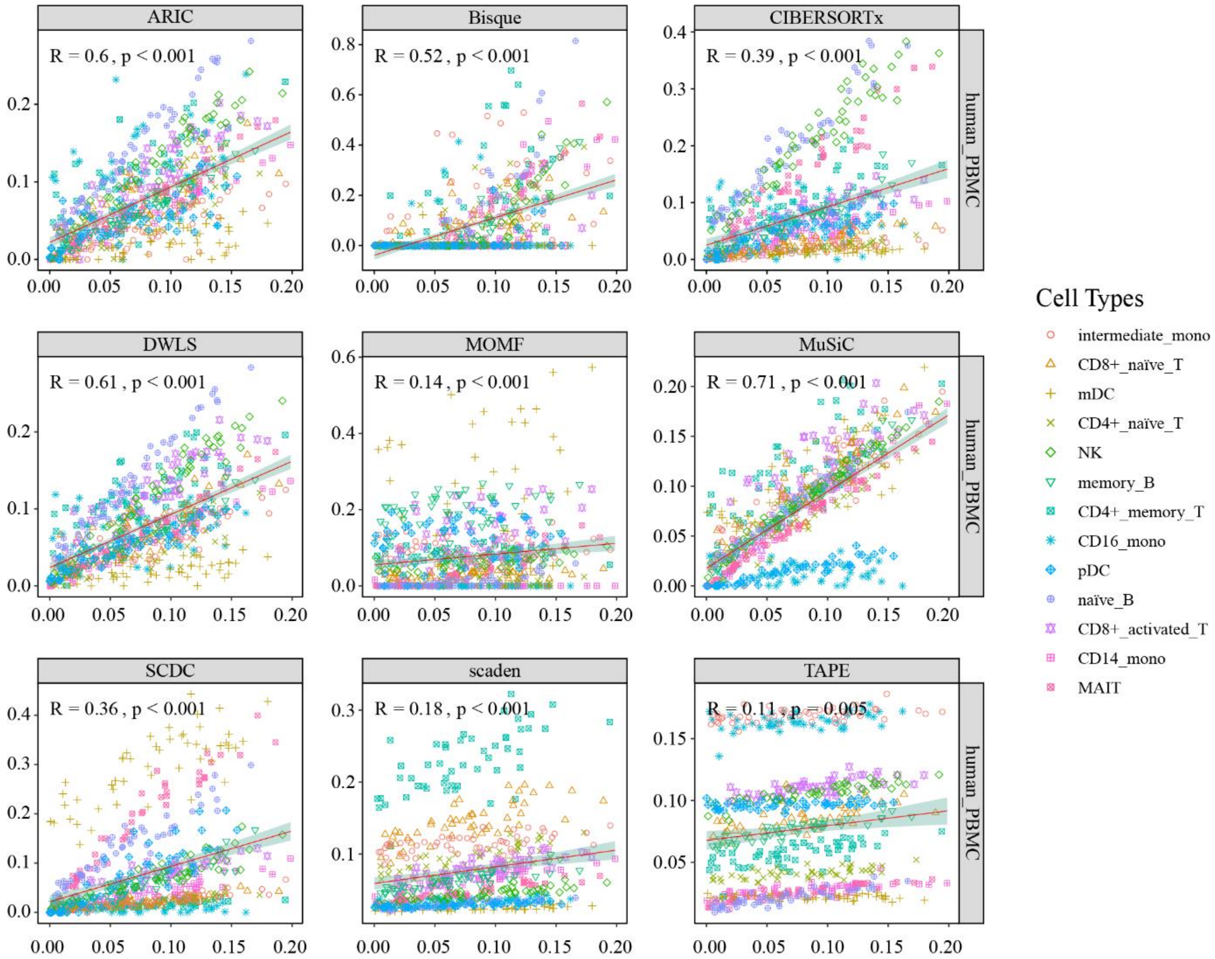


**Figure S8.** Deconvolution results for human_PBMC datasets. This figure corresponds to NL_0 in the figure S6, which means that noise is not added in this test. The datasets are annotated on the right side of the graph, and the names of each method are labelled above their respective graphs. The light green area represents the 95% confidence interval.

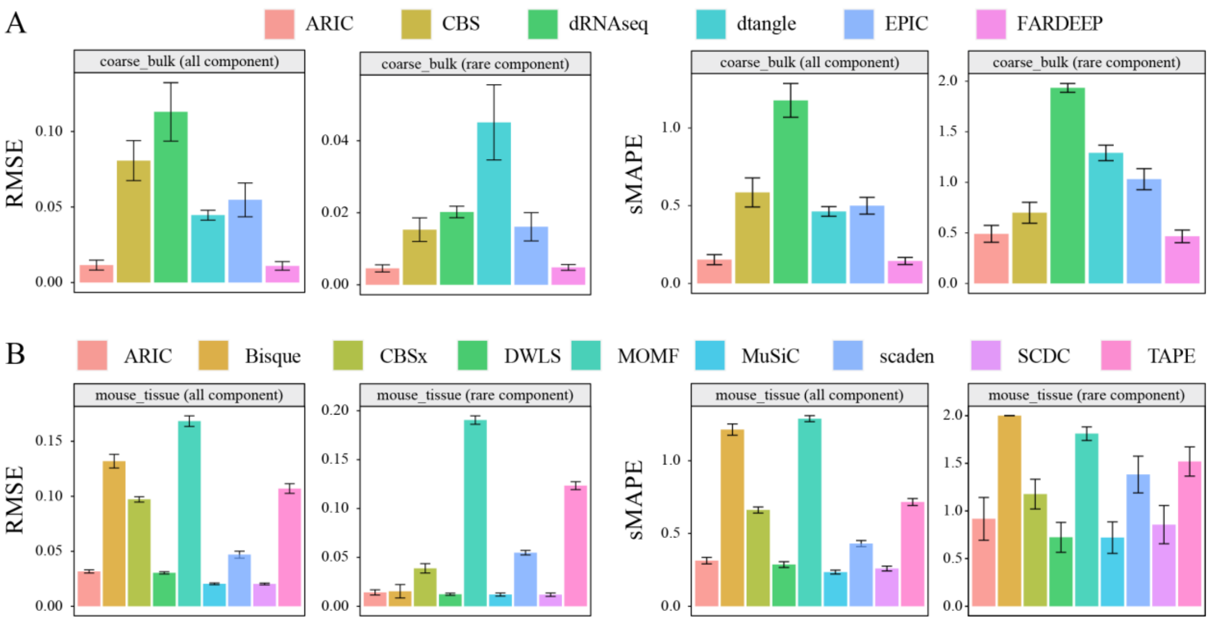


**Figure S9.** Impact of rare component on the deconvolution results using coarse_bulk and mouse_tissue datasets. All component as well as rare component are illustrated respectively.

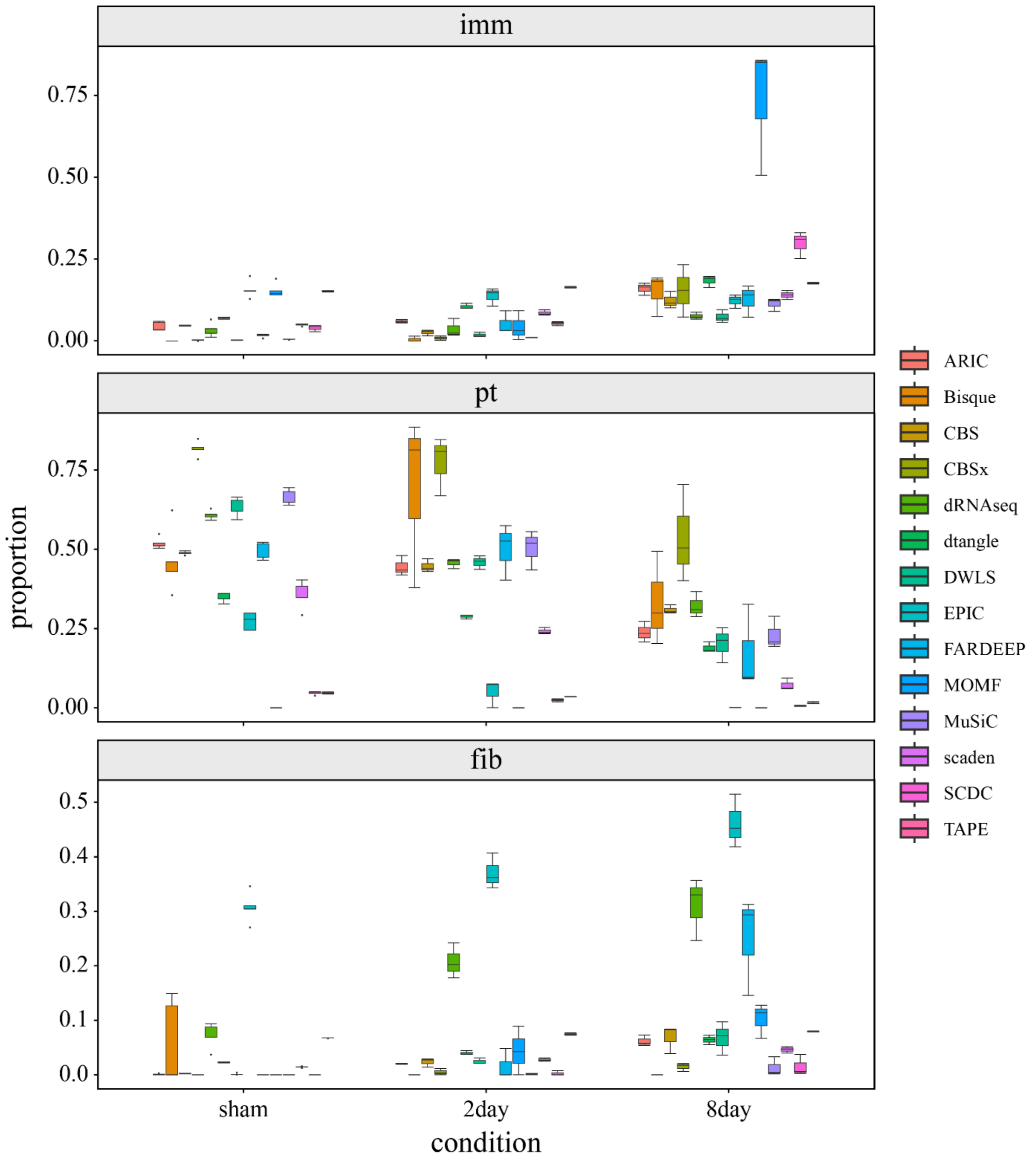


**Figure S10.** Proportions predicted by different methods for UUO model. The methods shown in the image are arranged in the same order as indicated in the legend on the right.
